# Successful axonal regeneration is driven by evolutionarily conserved metabolic reprogramming

**DOI:** 10.1101/2025.02.12.637870

**Authors:** Anyi Zhang, Steven Bergmans, Annelies Van Dyck, An Beckers, Lieve Moons, Luca Masin

## Abstract

Unlike mammals, zebrafish can regrow axons after injury and restore circuit function in the central nervous system (CNS). Mitochondria have been identified as key players in this process, but how different metabolic pathways work together to sustain regeneration remains unclear. Using RNA sequencing of adult zebrafish retinal ganglion cells after optic nerve crush injury, we found that oxidative phosphorylation is downregulated during axonal regrowth. Simultaneously, the thioredoxin antioxidant system was upregulated, likely to limit oxidative damage. Additionally, we observed an integrated upregulation of glycolysis and the pentose phosphate pathway during the initial regrowth phases, possibly to provide energy and supplying NADPH for biosynthesis and antioxidant responses. We show that this metabolic reprogramming is evolutionarily conserved, as a comparable one occurs in the pro-regenerative mammalian *Pten* and *Socs3* co-deletion model. Inhibiting glycolysis and thioredoxin in zebrafish impairs axonal regrowth, suggesting that targeting these pathways could enhance CNS regeneration in mammals.

## Introduction

The adult mammalian central nervous system (CNS) is extremely vulnerable to traumatic damage, due to its limited regenerative capacity. Injury leads to neuronal death (Varadarajan *et al*, 2022) and surviving neurons fail to regenerate axons due to intrinsic deficits in reprogramming gene expression and extrinsic inhibitory factors at the injury site (Fawcett, 2020). In mammals, regenerative capacity is progressively lost during development and is nearly absent after birth (Varadarajan *et al*, 2022). Conversely, adult teleost fish, such as zebrafish (*Danio rerio*), demonstrate remarkable CNS repair and functional recovery after injury (Lee-Liu *et al*, 2013; Bollaerts *et al*, 2017; Dhara *et al*, 2019). Their conserved genome and divergent responses to CNS injury provide a unique opportunity for comparative analysis to elucidate the molecular basis of successful CNS regeneration.

As part of the CNS, the adult zebrafish visual system is an established model for studying the regenerative response induced by optic nerve injury. Previous research using the optic nerve crush (ONC) injury model revealed a shift in mitochondrial density from the somatodendritic to the axonal compartment of the retinal ganglion cells (RGCs) during early axonal regeneration phases, followed by a return to baseline after reinnervation is complete (Beckers *et al*, 2023b). This suggests that mitochondria must relocate to support regrowth, consistent with findings in other regenerative models (Xu *et al*, 2017; Zhou *et al*, 2016; Han *et al*, 2016; Mar *et al*, 2014). Such relocation post-injury compensates for the inability of depolarized axonal mitochondria to supply energy (Petrova *et al*, 2021; Zhou *et al*, 2016). Unlike zebrafish, mammalian CNS neurons cannot redistribute mitochondria, leading to energy deficits, failure of axonal regrowth and eventual degeneration (Hopkins *et al*, 2021; Conforti *et al*, 2014). Evidence from the murine regeneration-competent model of conditional *Pten* and *Socs3* co-deletion in RGCs (Sun *et al*, 2011), indicates that axonal regrowth is accompanied by increased mitochondrial trafficking (Cartoni *et al*, 2017), but powered by local axonal glycolysis (Masin *et al*, 2024). This suggests that intraneuronal metabolic reprogramming might be required to at least initiate regeneration. This study aims to characterize the metabolic response of spontaneously regenerating adult zebrafish RGCs during injury-induced axonal regrowth to identify conserved mechanisms underlying regeneration and functional recovery.

To this end, we conducted bulk RNA sequencing on isolated adult zebrafish RGCs, collected from non-injured fish and at various timepoints post-ONC, covering the complete regeneration time window. Early axonal regrowth phases showed downregulation of oxidative phosphorylation (OXPHOS) genes and upregulation of glycolysis and the thioredoxin antioxidant system, likely supporting regeneration by providing energy and neutralizing reactive oxygen species (ROS) generated by damaged mitochondria. Additionally, the pentose phosphate pathway (PPP) and cholesterol biosynthesis were upregulated, supplying essential building blocks for axonal extension. Significant overlap was found between the upregulated pathways in our data and those previously reported in the murine *Pten* and *Socs3* co-deletion model (Jacobi *et al*, 2022). Validation using ex vivo retinal explants and in vivo approaches showed that inhibiting glycolysis or the thioredoxin system impaired axonal regrowth and delayed optic tectum reinnervation, suggesting these pathways could be targeted to enhance mammalian CNS regeneration.

## Results

### Transcriptomic profiling of spontaneously regenerating adult zebrafish RGCs following ONC injury

To investigate the transcriptional programs and metabolic gene signatures associated with axonal regeneration and functional recovery post-ONC, we performed bulk RNA sequencing of fluorescently sorted zebrafish RGCs (Fig. 1A; Fig. S1). Unlike whole-retina sequencing studies (Dhara *et al*, 2019), this approach captures RGC-specific expression, otherwise masked by other cell types. To achieve this, we isolated RGCs using *Tg(isl2b:eGFP)*^*zc7Tg*^ adult fish, which label most RGC subtypes (Kölsch *et al*, 2021). To capture the full regeneration timeline, we sampled 6 conditions: a non-injured naive control and 5 post-injury timepoints - 1, 3, 6, 10, and 14 days post-ONC injury (dpi), chosen based on in-house (Beckers *et al*, 2019) and published (Dhara *et al*, 2019) regeneration stage characterizations. These represent axonal regrowth initiation (1dpi), growth toward the midline at the optic chiasm (3dpi), optic tectum reinnervation (6dpi), target contact and synaptic refinement (10dpi), completion of tectal reinnervation with initial visual recovery (14dpi) (Fig. 1B).

**Fig. 1.**
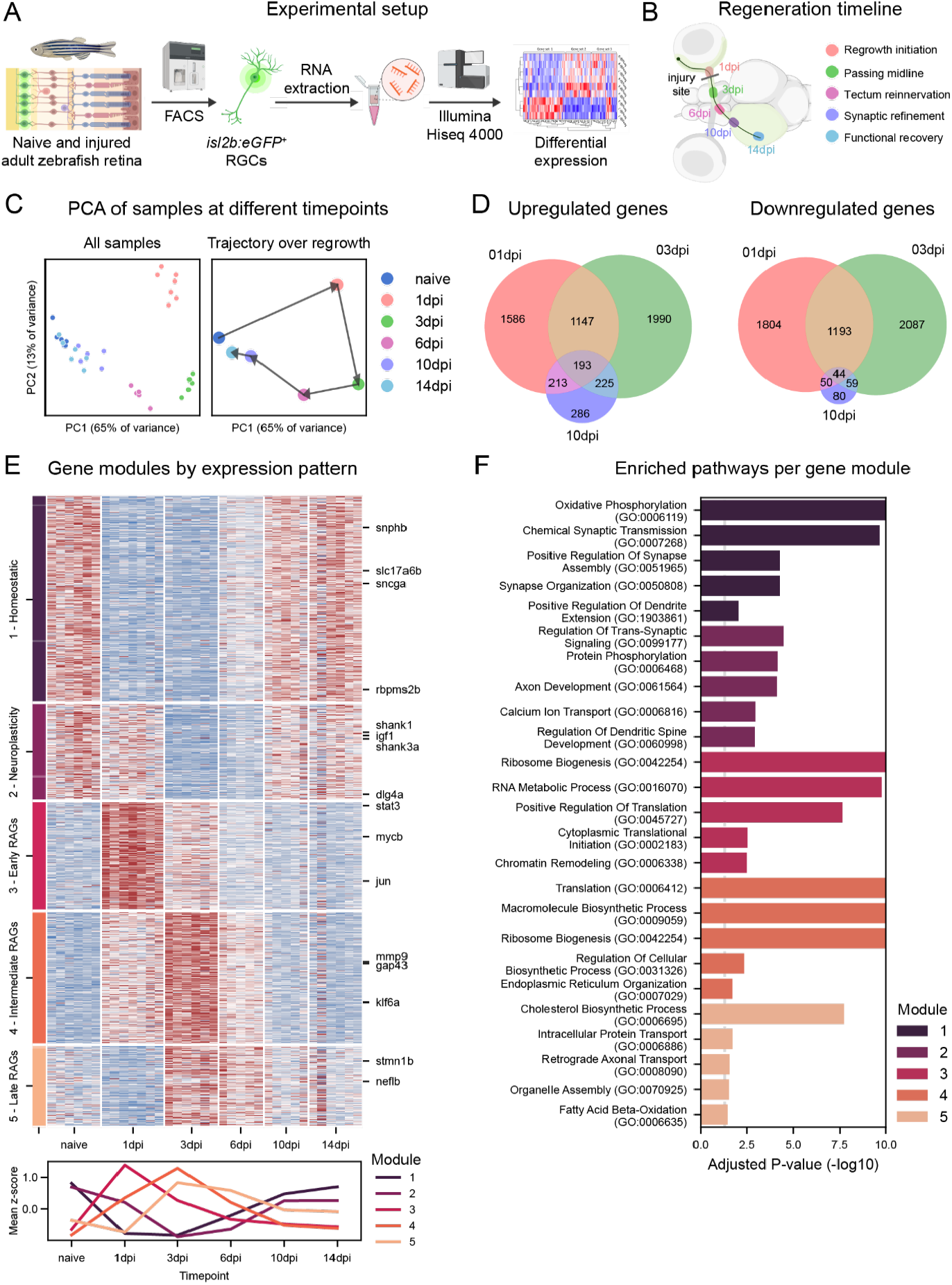
Axonal regeneration of adult zebrafish RGCs is driven by profound transcriptomic reprogramming. (A) Experimental setup for bulk RNA sequencing of FAC-sorted adult zebrafish RGCs, sampled from naive and injured (including 1, 3, 6, 10, and 14dpi) *Tg(isl2b:eGFP)*^*zc7Tg*^ fish. (B) Schematic overview of axonal regeneration phases following ONC injury. At 1dpi, axonal regrowth begins. Regenerating axons extend toward the midline at the optic chiasm by 3dpi and begin to reinnervate the optic tectum by 6dpi. By 10dpi, regenerating axons reconnect with target neurons with visual function starting to recover by 14dpi. (C) Principal component analysis (PCA) of naive samples and samples from different timepoints post-injury (n=5-7). The trajectory over the regrowth process indicated that profound transcriptomic reprogramming occurred as early as 1dpi, and as regeneration progresses, gene expression gradually returned to the naive state. (D) Venn diagram depicting the upregulated (left) and downregulated (right) genes across 1 (red), 3 (green), and 10dpi (purple). While a small portion of genes showed consistent differential expression trends across all three timepoints, the majority of transcripts were uniquely regulated, indicative of distinct axonal regeneration phases. (E) Heatmap showing the expression of DEGs across time, K-means clustered into distinct gene modules (top). The transcriptomic profile of regenerating RGCs evolved through 5 distinct stages, characteristic of different phases of axonal regrowth and visual function restoration. Aggregate expression profiles of each gene modules were represented at the bottom. (F) Pathway overrepresentation analysis of genes in each module. Early modules showed enrichment for homeostasis-related pathways, while later ones were enriched for translation and biosynthetic functions. Abbreviation: dpi (day post injury), DEGs (differentially expressed genes), FACS (fluorescence-activated cell sorting), ONC (optic nerve crush). Figure containing assets from BioRender.com.

After quality control, read mapping, gene filtering, and batch correction, we performed principal component analysis (PCA) on biological replicates across all timepoints. The replicates of each timepoint clustered closely, forming a trajectory that captured the progression of regeneration (Fig. 1C). Significant transcriptomic changes emerged as early as 1dpi, with further changes occurring at 3 and 6dpi, as shown by their distinct segregation on the PCA plot. Subsequently, as regeneration progressed, the transcriptomic profile began reverting to a naive-like state (Fig. 1C). Differential gene expression analysis revealed thousands of differentially expressed genes (DEGs) (log2FC > 1, FDR < 0.05), many of which were timepoint-specific, while a small subset overlapped across adjacent timepoints (Fig. 1D). These results underscore the extensive and temporally orchestrated transcriptomic reprogramming essential for successful axonal regeneration.

### Distinct regeneration phases are characterized by unique gene modules

To obtain a broad picture of the transcriptomic evolution across the regeneration timeline, we used K-means clustering to group the DEGs of all timepoints based on their shared expression patterns, identifying five distinct modules with similar profiles (Fig. 1E; Table.S1). Pathway overrepresentation analysis with GSEApy provided insights into the cellular processes associated with the genes of each module (Fig. 1F). The first one contained homeostatic genes characteristic of non-injured RGCs, which were downregulated immediately at 1dpi and gradually restored after 6dpi (Fig. 1E). Pathway analysis showed that genes in this module were significantly enriched for OXPHOS, synaptic function and organization, and dendrite extension (Fig. 1F). Notably, this module included general RGC markers, such as *rbpms2b* (RNA-binding protein with multiple splicing), *sncgb* (synuclein gamma) and *slc17a6b* (vglut2, vesicular glutamate transporter 2), as well as *snphb* (syntaphilin), a mitochondrial docking protein whose deletion promotes axonal regeneration in the mouse CNS (Han *et al*, 2020; Zhou *et al*, 2016). The second module comprised of neuroplasticity-related transcripts that largely retained expression during the injury response phase (1dpi) but were downregulated as regrowth began (3dpi), eventually returning to baseline (Fig. 1E). Pathway analysis revealed significant enrichment in synaptic regulation, dendritic remodeling, and axonogenesis (Fig. 1F). This module included key postsynaptic genes, such as *dlg4a* and *dlg4b* (*PSD-95, postsynaptic density protein 95*) and *shank1* and *shank3a* (SH3 and multiple ankyrin repeat domains proteins), indicating reduced RGC postsynaptic assembly during axonal elongation, consistent with prior findings (Beckers *et al*, 2019). Additionally, we identified genes linked to RGC survival, such as *igf1* (insulin-like growth factor 1) and *bdnf* (brain-derived neurotrophic factor) (Tapia *et al*, 2022). In the third gene module (early regeneration-associated genes, RAGs), transcripts were rapidly upregulated post-injury, sustained during early regeneration, and later returned to baseline (Fig. 1E). This module was enriched in RAGs, including *stat3* (signal transducer and activator of transcription 3) (Leibinger *et al*, 2013; Luo *et al*, 2016), *mycb* (c-Myc, myelocytomatosis oncogene) (Belin *et al*, 2015), and *jun* (Jun proto-oncogene) (Mason *et al*, 2021). Pathway analysis identified enrichment for translation-related processes (Fig. 1F), suggesting extensive transcriptional reprogramming in RGCs. The fourth module (intermediate RAGs) contained transcripts that peaked during axonal elongation at 3dpi, remained upregulated at 6dpi, and returned to baseline by 10dpi (Fig. 1E). This module was also enriched for translation and biosynthesis (Fig. 1F), and additionally, it included genes related to axonal regrowth, such as *gap43* (growth-associated protein-43) (Wang *et al*, 2022), and pro-regeneration factors, like *klf6a* (Kruppel like factor 6a) (Kramer *et al*, 2021) and *mmp9* (matrix metallopeptidase 9) (Andries *et al*, 2016). The final module (late RAGs) consisted of transcripts upregulated at 3 and 6dpi (Fig. 1E). Pathway analysis highlighted a strong enrichment only for cholesterol biosynthesis (Fig. 1F), likely due to the small gene set. Nonetheless, we identified genes related to microtubules, including *stmn1b* (stathmin 1b) (Gagliardi *et al*, 2022) and *neflb* (neurofilament light chain b) (Wang *et al*, 2012), which are implicated in axonal regeneration. Together, these results demonstrate that successful axonal regeneration is driven by a dynamic and phase-specific evolution of transcriptional landscapes.

### Pathway analysis reveals unique signatures of metabolic reprogramming during axonal regeneration

To investigate the differentially expressed pathways driving each regenerative phase, we performed gene set enrichment analysis (GSEA) on the DEGs of each timepoint. To provide a comprehensive and consensual view, we utilized three different databases/tools: GSEA on GO Biological Processes and Reactome databases (for its rich annotation of metabolic pathways) with GSEApy (Fang *et al*, 2023), along with pathway analysis using Ingenuity Pathway Analysis (IPA) (for its greater annotation of higher-order pathways) (Fig. 2A). Comparing the naive condition with early regeneration timepoints (1dpi to 6dpi), at least two out of three databases agreed on increased pathway enrichment for Akt activation downstream of PIP3, translation, ribosome biosynthesis, and interleukin-1 signaling, all pathways known to play a central role in axonal regrowth (Schaeffer *et al*, 2023; Huang *et al*, 2021; Akram *et al*, 2022; Williams *et al*, 2020). All databases highlighted the downregulation of synaptic organization/transmission and OXPHOS between 1dpi and 6dpi (Fig. 2B-D) and the enrichment of cholesterol synthesis pathways between 3dpi and 6dpi (Fig. 2B-D). GO uniquely showed downregulated dendrite spine development, peaking at 3dpi (Fig. 2B), aligning with previous findings of dendritic shrinkage in early phases of RGC axonal regeneration (Beckers *et al*, 2019). GSEA on Reactome and IPA detected significant upregulation of glycolysis/glucose metabolism between 1dpi and 3dpi (Fig. 2C-D), which might serve as energy source to initiate regeneration. Uniquely, IPA identified a significant and strong upregulation of NRF2-mediated oxidative stress response between 1dpi and 10dpi, and, specifically at 3dpi, a significant upregulation of PPP, which also plays a central role in antioxidant responses (Fig. 2C) (TeSlaa *et al*, 2023). Further, IPA identified sustained microautophagy upregulation throughout regeneration, consistent with previous findings showing its induction in adult zebrafish RGCs after ONC (Beckers *et al*, 2021) (Fig. 2D). Taken together, these enrichment analyses highlighted multiple signaling pathways previously reported to be critical for regeneration, including mTOR, STAT, and inflammation. Regarding metabolism, all three analyses suggested a strong downregulation of OXPHOS. Notably, IPA highlighted the NRF2 pathway of antioxidant response and the PPP, while Reactome pointed out glycolysis. However, since the rate of most metabolic pathways is governed by specific rate-limiting enzymes, a closer examination of these at the single gene level is necessary to more confidently infer metabolic activities.

**Fig. 2.**
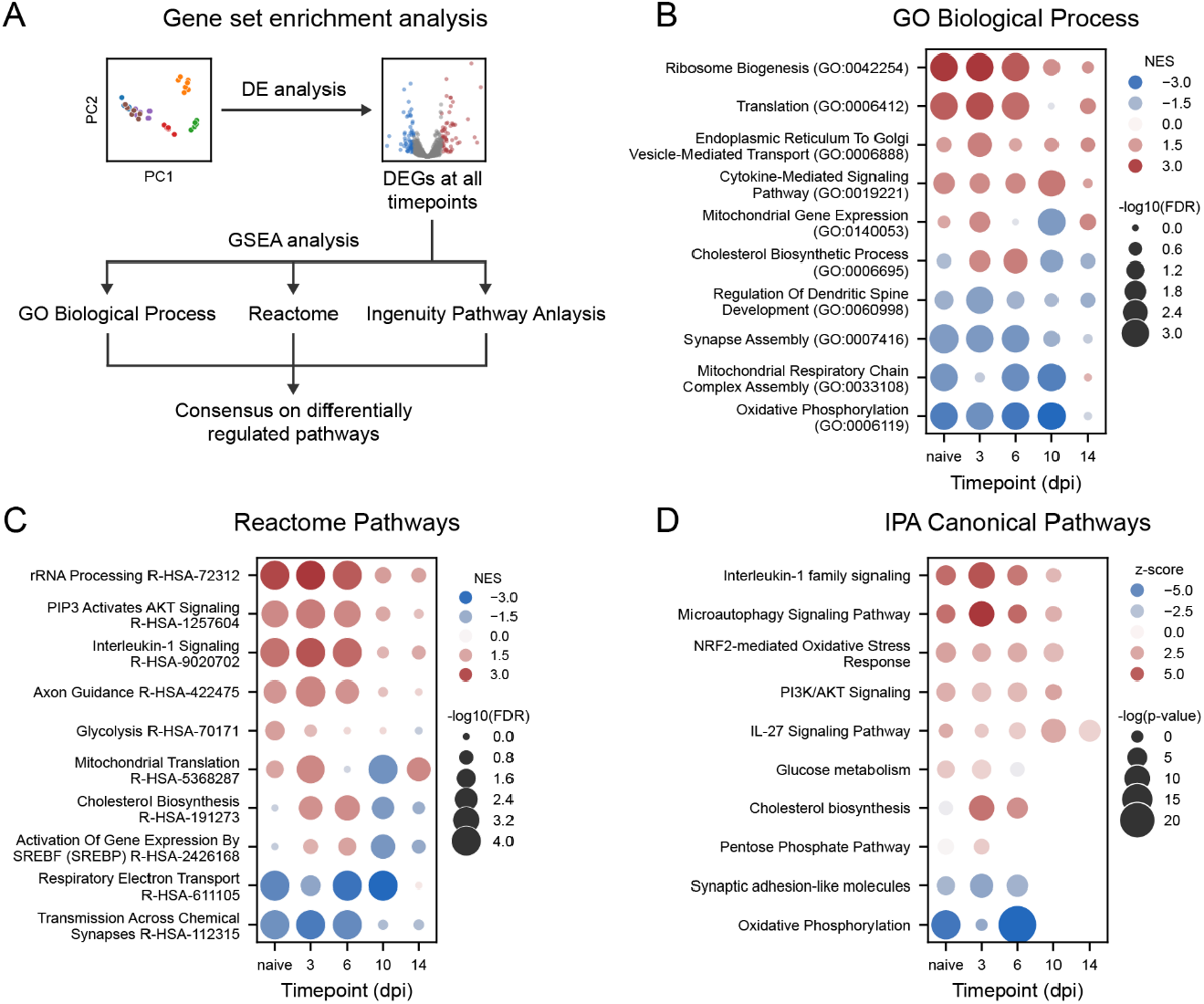
GSEA reveals pathway signatures associated with distinct axonal regeneration phases. (A) GO Biological Process, Reactome, and Ingenuity Pathway Analysis (IPA) were used to provide a comprehensive and consensual overview of the pathways differentially expressed during various regeneration stages. (B) GSEA using GO Biological Process highlighted increased translation activity during early axonal regrowth phases (1 to 6dpi) and elevated cholesterol biosynthesis during axonal elongation (3 to 6dpi). It also revealed downregulated dendrite-related GO terms and OXPHOS. (C) GSEA using Reactome pathways similarly identified augmented cholesterol biosynthesis and reduced OXPHOS. Additionally, it showed enhanced glycolysis at early regeneration stages. (D) GSEA with IPA confirmed upregulation of cholesterol biosynthesis and downregulated OXPHOS. Furthermore, it identified elevated antioxidant responses and activation of the pentose phosphate pathway during early regeneration stages. Abbreviation: DE (differential expression), DEG (differentially expressed genes), dpi (day post injury), GO (gene ontology), GSEA (gene set enrichment analysis, IPA (Ingenuity Pathway Analysis), PC (principle component), NES (normalized enrichment score), FDR (false discovery rate).

### Injury-induced axonal regrowth in zebrafish RGCs is associated with a profound transcriptional reprogramming of metabolism

To assess metabolic reprogramming, we analyzed the differential expression of key rate-limiting enzymes of five metabolic pathways which were differentially expressed at the early stages of axonal regeneration: 1) glycolysis, 2) mitochondrial-related pathways, 3) the PPP, 4) the mevalonate and cholesterol pathways, and 5) antioxidant pathways. Specifically, we examined the expression at 1 and 3dpi, corresponding to the initiation and early elongation phases of axonal regrowth (Fig. 3). Expression was validated in situ, via hybridization chain reaction (HCR) or RNAscope (Fig. 4). This analysis focused on the RGC layer, though expression changes could also be observed in the inner plexiform and inner nuclear layers (Fig. 4). While differential expression in other retinal cell types may contribute to RGC axonal regeneration, further investigation is beyond the scope of this study.

**Fig. 3.**
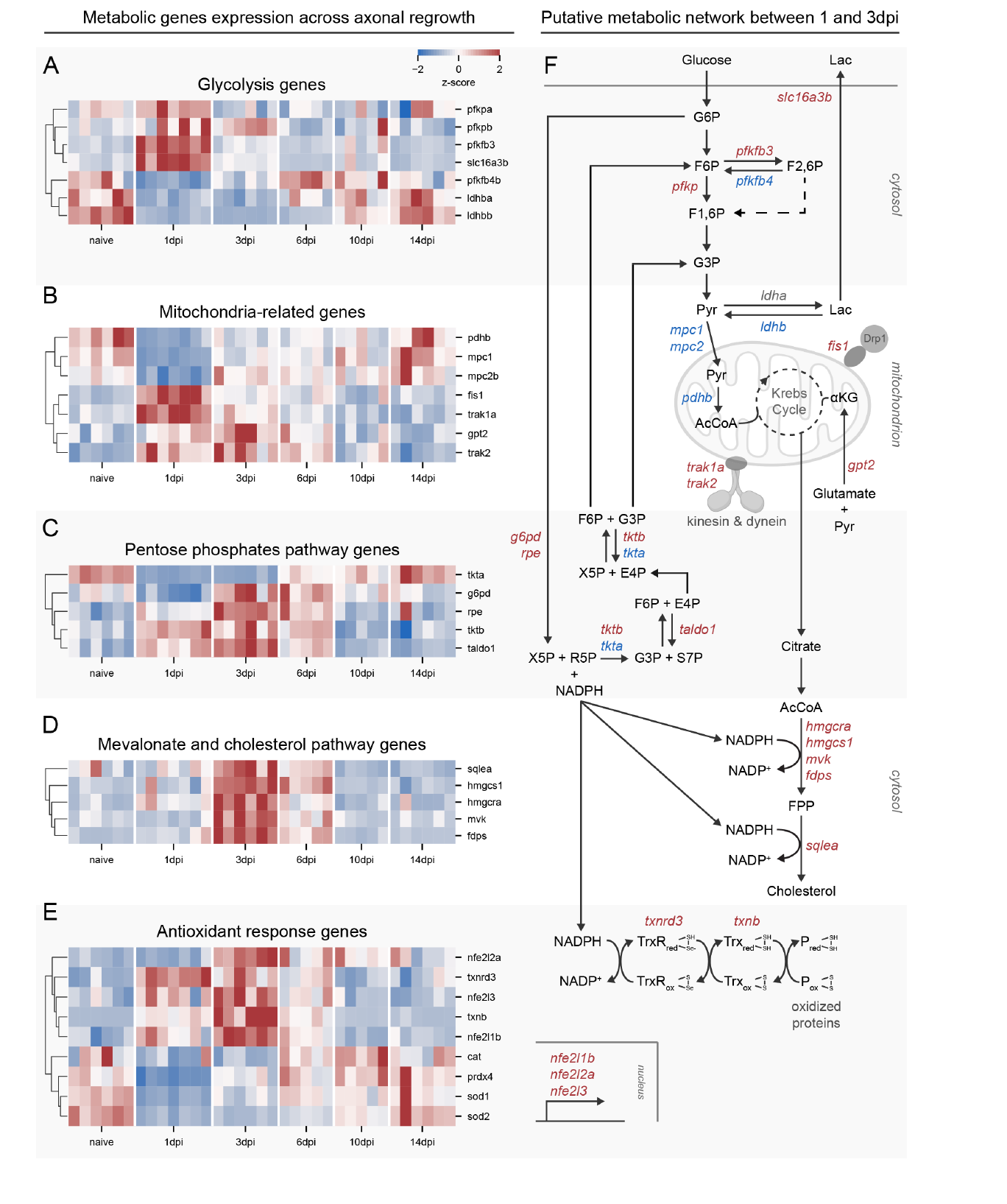
Axonal regrowth is associated with a profound reprogramming of carbon metabolism and antioxidant response. (A) Heatmap showing the z-scored expression of glycolytic genes. At 1 day post injury (dpi), the main rate-limiting genes were upregulated, suggesting an increased activity of the pathway during early phases of axonal regeneration. (B) Z-scored expression of genes related to mitochondrial metabolism and dynamics. At 1dpi and to a lesser extent until 6dpi, genes related to the import of pyruvate into the mitochondria for OXPHOS were downregulated. In contrast, genes related to mitochondrial fission and transport along microtubules were upregulated. (C) Z-scored expression of genes belonging to PPP. Genes of the non-oxidative branch were upregulated at 1dpi, followed by an upregulation of the whole pathway at 3 and 6dpi. (D) Z-scored expression of genes related to the synthesis of mevalonate and cholesterol. Most genes of the two pathways showed increased expression between 3 and 6dpi. (E) Z-scored expression of genes related to cellular antioxidant defence. The transcription factors regulating antioxidant response showed upregulation between 1 and 3dpi. Downstream, the genes of the thioredoxin system were upregulated, while superoxide dismutase and catalase were downregulated. (F) Schematic representation of the integrated metabolic network between 1 and 3dpi, inferred from the transcriptomic data. Early after injury, glycolysis is upregulated, and OXPHOS is downregulated. Branching from glycolysis, the PPP generates NADPH, essential for the synthesis of mevalonate derivatives and cholesterol, as well as for limiting oxidative damage. Gene expression profiles are presented along with hierarchical clustering of their expression profile. Abbreviations: dpi (day post injury), G6P (glucose-6-phosphate), F6P (fructose-6-phosphate), F1,6P (fructose-1,6-bisphosphate), F2,6P (fructose-2,6-bisphosphate), G3P (glyceraldehyde-3-phosphate), X5P (xylulose-5-phosphate), R5P (ribose-5-phosphate), S7P (sedoheptulose-7-phosphate), E4P (erythrose-4-phosphate), Pyr (pyruvate), Lac (lactate), αKG (α-ketoglutarate), AcCoA (Acetyl-CoA), FPP (farnesylpyrophosphate).

**Fig. 4.**
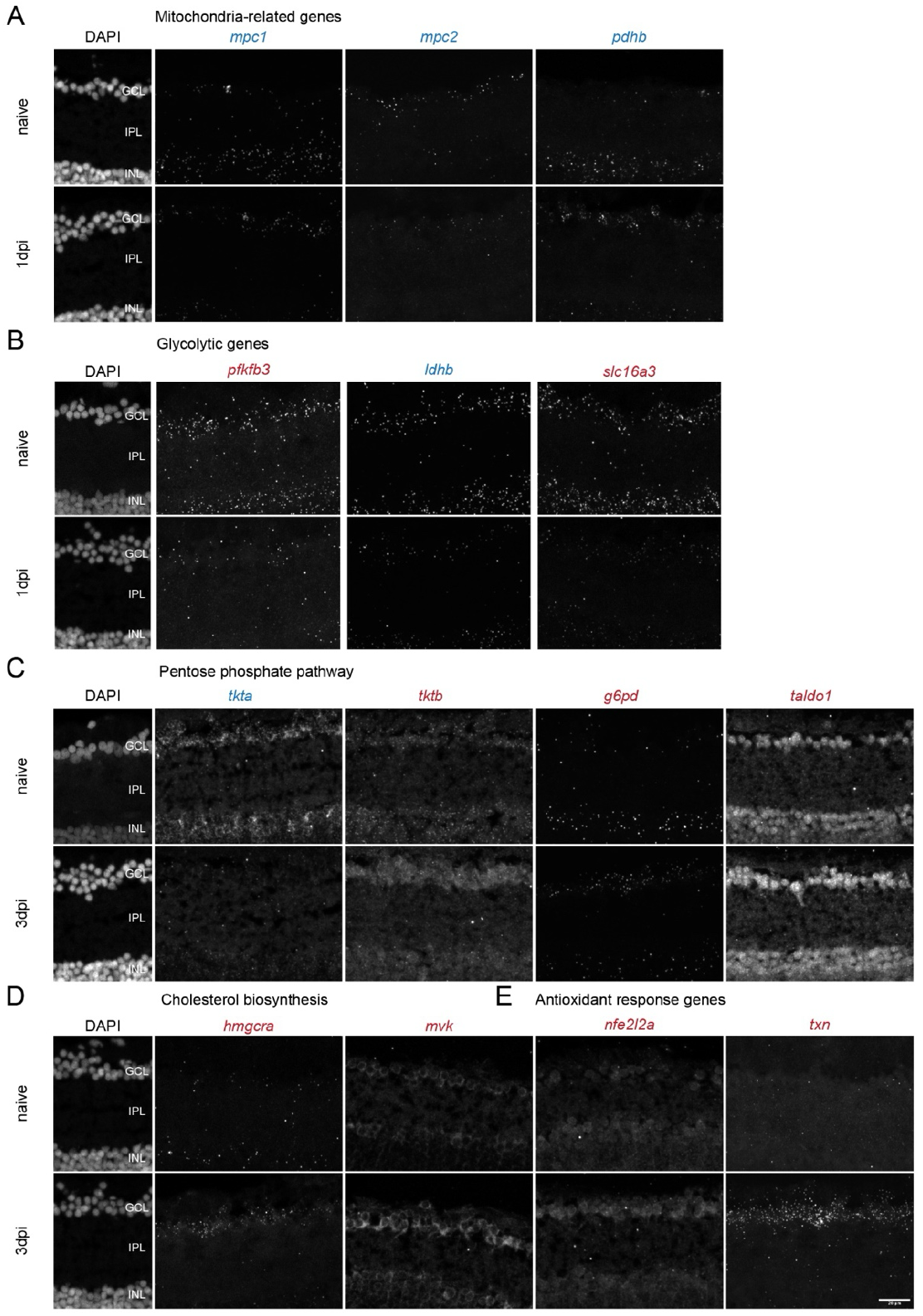
The differential mRNA expression of key metabolic genes was confirmed using in situ hybridization. (A) At 1dpi, mRNA expression for the glycolytic genes *pfkfb3* and *slc16a3* was upregulated, while *ldhb* was downregulated in the GCL. (B) mRNA expression for mitochondria-related genes, including the mitochondrial pyruvate carriers *mpc1* and *mpc2*, and the Krebs cycle gene *pdhb*, was downregulated at 1dpi in the GCL. (C) mRNA expression of PPP-related genes was differentially regulated during early axonal regrowth (1dpi to 3dpi). *tkta* was downregulated, while *g6pd, tktb*, and *taldo1* were upregulated at 3dpi in the GCL. (D) Expression of mRNA for genes related to cholesterol biosynthesis, including *hmgcra* and *mvk*, was augmented at 3dpi in the GCL. (E) The expression of mRNA for genes related to the antioxidant response, including *nfe2l2a* and *txn*, was upregulated at 3dpi in the GCL. Expression of mRNA for *tkta, tktb, taldo1, mvk*, and *nfe2l2a* was visualized using HCR. The remaining genes were labeled using RNAscope. All genes that showed upregulated mRNA expression are annotated in red, the downregulated ones in blue. AB wild-type fish were used for this experiment. n=4. Scale bar 20μm. Abbreviation: dpi (days post-injury), GCL (ganglion cell layer), INL (inner nuclear layer), IPL (inner plexiform layer).

We first analyzed the differential expression of key glycolytic enzymes (Fig. 3A) and observed a slight upregulation at 1dpi of both *pfkpa* and *pfkpb* (PFKP, phosphofructokinase, platelet isoform) the main rate-limiting enzyme in glycolysis (Fig. 3A, F; Table.S2). Furthermore, a strong and significant upregulation of *pfkfb3* (Fig. 3A, F; Fig. 4A) and a significant downregulation of *pfkfb4* (6-phosphofructo-2-kinase/fructose-2,6-biphosphatase 3 and 4, respectively) (Fig. 3A, F) was observed at 1dpi. *pfkfb3* favors the production and *pfkfb4* the breakdown of fructose-2,6-bisphosphate, a strong allosteric activator of PFKP (Fig. 3F). Together, this suggests an increased rate of glycolysis via the allosteric activation of PFK activity. Additionally, *slc16a3b* (MCT4, monocarboxylate transporter 4), the primary transporter for lactate export, was upregulated at 1dpi (Fig. 3A, F; Fig. 4A), while *ldhb* (lactate dehydrogenase b), which converts lactate back into pyruvate, was downregulated at 1dpi (Fig. 3A, F; Fig. 4A). These expression patterns suggest that RGCs are primarily glycolytic during the early regrowth phases and may produce and export lactate, consistent with the Warburg effect (Liberti & Locasale, 2016; Heiden *et al*, 2009). This hypothesis is reinforced by the detected downregulation of genes linking glycolysis to OXPHOS at 1dpi, including *mpc1* and *mpc2* (MPC, mitochondrial pyruvate carrier 1 and 2) (Fig. 3B, F; Fig. 4B), which shuttle pyruvate into the mitochondrion, and *pdhb*, a subunit of pyruvate dehydrogenase that converts pyruvate into acetyl-CoA to feed the Krebs cycle (Fig. 3B, F; Fig. 4B). In contrast, *gpt2* (GPT2, glutamic pyruvate transaminase 2) was upregulated between 1 and 3dpi (Fig. 3B). In muscle, this enzyme catalyzes the conversion of pyruvate and glutamate into alanine and α-ketoglutarate, with the latter feeding the Krebs cycle (Fig. 3F). In injured neurons, it may help utilize excess glutamate as an energy source and reduce excitotoxicity (Divakaruni *et al*, 2017). Regarding mitochondrial dynamics, we detected an upregulation of *trak1a* and *trak2*, which are adaptor protein connecting mitochondria to dynein and kinesin motors, indicating increased mitochondrial transport (Fig. 3B, F). *fis1*, a mitochondrial receptor of Drp1 mediating fission, was also elevated at 1dpi (Fig. 3B, F) (Giacomello *et al*, 2020). Together these findings suggest reduced OXPHOS involvement in early regeneration, with RGCs relying more on glycolysis for energy production.

Branching from glycolysis, at 3dpi we observed a minor upregulation of *g6pd* (glucose-6-phosphate dehydrogenase) (Fig. 3C, F; Fig. 4C; Table.S2), the rate-limiting and first enzyme of the oxidative branch of the PPP, and of the downstream enzyme *rpe* (ribulose-5-phosphate-3-epimerase) (Fig. 3C, F). The oxidative PPP (oxPPP) replenishes NADPH, which is crucial for antioxidant responses and biosynthetic pathways, while the non-oxidative PPP (non-oxPPP) recycles the end-product of the oxidative branch into glycolysis intermediates for energy production or precursors for nucleotide biosynthesis (Fig. 3F). Differential expression of genes of the non-oxPPP began at 1dpi but was sustained until 6dpi. Specifically, we detected a significant increase in expression of *taldo1* (transaldolase 1) and a switch in expression from *tkta* to *tktb* (transketolase a and b) (Fig. 3C, F; Fig. 4C). In summary, this upregulation of the PPP is likely required to preserve NADPH levels while also maintaining a high rate of glycolysis.

NADPH is a major cofactor for lipid biosynthesis. We found an upregulation of the majority of enzymes of mevalonate pathway, including hmgcra (3-hydroxy-3-methylglutaryl-CoA reductase a), the rate-limiting enzyme, and the downstream mvk (mevalonate kinase) (Fig. 3D, F; Fig. 4D). Additionally, we detected increased expression of sqle (squalene oxidase), the rate-limiting enzyme in the downstream synthesis of cholesterol (Fig. 3D, F). The increase in cholesterol production could be required to modulate the fluidity of the plasma membrane for active growth cone formation during axonal extension and/or the organization of membrane receptors on lipid rafts (Korade & Kenworthy, 2008; Roselló-Busquets et al, 2019). Finally, NADPH serves as an electron donor in antioxidant detoxification mechanisms (An et al, 2024) (Fig. 3E-F). A specific and strong upregulation of the thioredoxin system was noted between 1dpi and 3dpi, including a marked upregulation of txnb (thioredoxin b) (Fig. 3E-F; Fig. 4E), which detoxifies oxidized proteins, rather than more canonical mechanisms targeting directly hydrogen peroxide (H_2_O_2_) or superoxide. The transcription factors of NRF signaling (nfe2l1b, nfe2l2a, and nfe2l3), which are upstream and likely responsible for this antioxidant response, were co-upregulated between 1dpi and 3dpi (Fig. 3E-F; Fig. 4E). In summary, the transcriptomic signature of RGCs during the early phases of axonal regeneration suggests a switch from a mitochondrial to glycolytic metabolism. The increase in glycolysis is accompanied by an upregulation of the related PPP, which is thought to provide NADPH for antioxidant response and lipid biosynthesis.

### The core metabolic reprogramming is conserved between zebrafish and a murine model of induced axonal regrowth

Recent evidence shows that, in regeneration-competent *Pten* and *Socs3* co-deleted RGCs, injury-induced axonal regrowth is fueled by local glycolysis within the axon (Masin *et al*, 2024). Thus, we next explored how transcriptional reprogramming in adult zebrafish compares to this murine model of induced axonal regeneration (conditional co-deletion of *Pten* and *Socs3* in RGCs, referred to here as KO). As in mice target reinnervation and functional recovery do not occur, we focused the comparison on the early phase of regrowth initiation. To this end, we compared the DEGs at 1dpi in our zebrafish dataset with the DEGs of KO RGCs at the earliest timepoint post-ONC (2dpi) in a murine scRNAseq dataset (Jacobi *et al*, 2022), reanalyzed as pseudo-bulk (Masin *et al*, 2024) (Fig. 5A). For both, the genes were mapped to human orthologues. The two species were found to share 419 genes with the same expression pattern, i.e. significantly up- or down-regulated early upon injury and 127 genes with opposite expression patterns. Nevertheless, many genes were uniquely differentially expressed in each organism (Fig. 5B). Pathway overrepresentation analysis revealed that the shared genes were significantly enriched for regulation of axonogenesis, cellular response to calcium, synapse regulation, dendrite extension and the KEAP-NFE2L2 (NRF2) pathway of stress response (Fig. 5B), while genes with opposite confidently mapped only to cholesterol biosynthesis. Overall, among the genes identified in our previous analysis in zebrafish, the ones associated with the NRF2 pathway, synapse assembly and dendrite extension, emerged as the most conserved among regenerative vertebrate models.

**Fig. 5.**
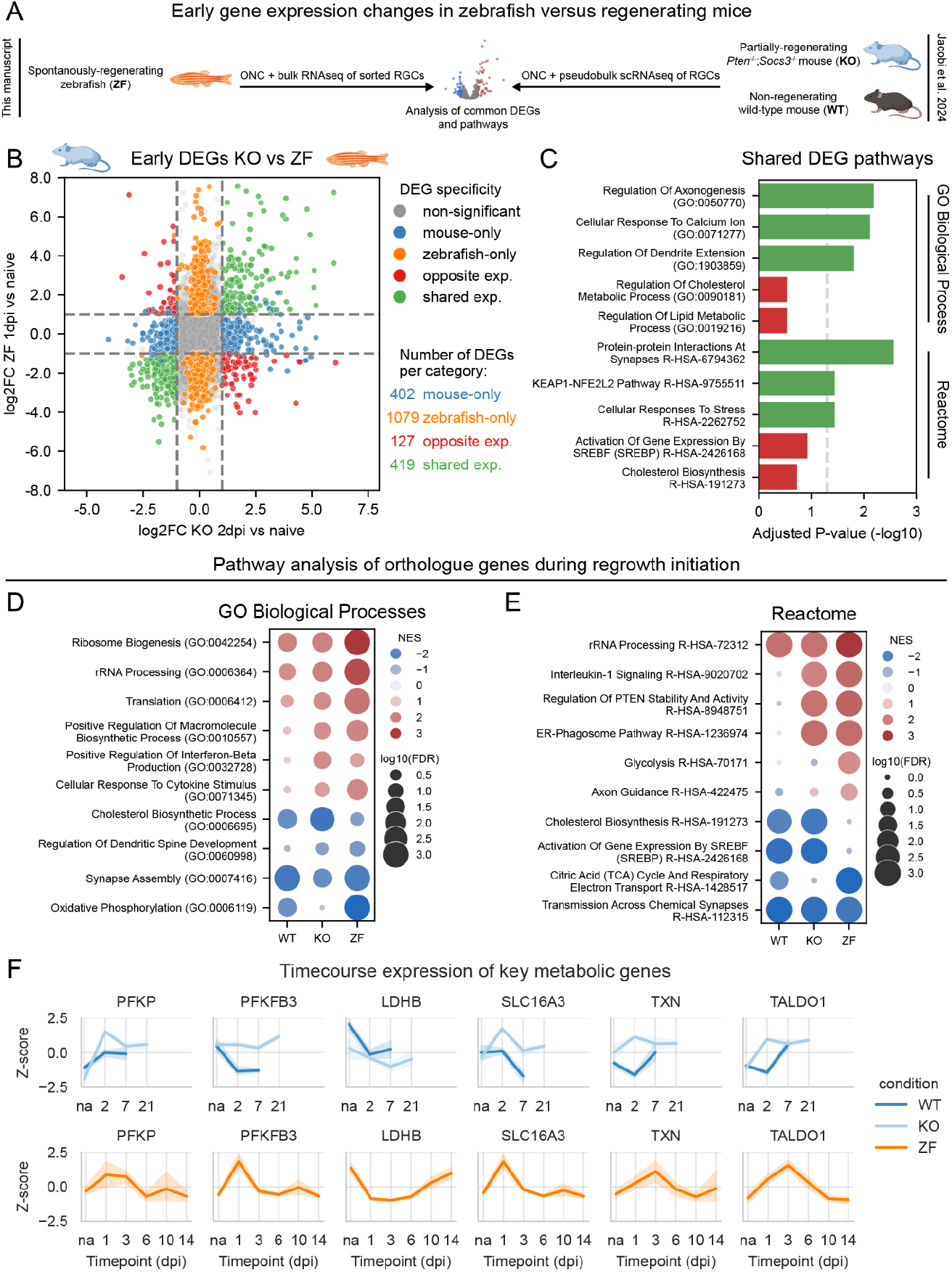
The upregulation of glycolysis and thioredoxin is an evolutionarily conserved feature of injury-induced axonal regeneration. (A) Zebrafish (ZF) RGCs transcriptomic data from this study was compared to Jacobi et al’s dataset of wild-type (WT) and *Pten and Socs3 co-deleted RGCs* (KO) post-ONC (Jacobi *et al*, 2022). To compare early axonal regeneration, we picked the earliest timepoint in both datasets, 1dpi and 2dpi, respectively. For cross-species DEGs comparison, genes were mapped to their human orthologues. (B) Scatterplot showing the differential expression of genes between zebrafish and KO mice. Genes are colored by expression pattern: shared (green), opposite (red), uniquely significant in zebrafish (orange), uniquely significant in KO mice (blue), and non-significant (grey). (C) Pathway overrepresentation analysis of genes with shared (green) and opposite (red) expression patterns. GO BP and Reactome analysis showed shared genes involved in axonogenesis, antioxidant response, and dendrite remodeling, while opposing genes were linked to cholesterol biosynthesis. (D) GSEA analysis of GO BP for orthologue DEGs from WT mice, KO mice, and zebrafish showed upregulation of ribosome-related pathways and downregulation of synapse assembly. KO mice and zebrafish both exhibited activation of biosynthesis- and inflammation-related pathways, and downregulation of dendrite spine development. (E) GSEA analysis of Reactome pathways for orthologue DEGs from WT mice, KO mice, and zebrafish showed upregulation of ribosome biogenesis and downregulation of synaptic transmission. KO mice and zebrafish activated inflammation- and ER-phagosome-related pathways. Both mouse genotypes uniquely downregulated cholesterol biosynthesis, while zebrafish showed a stronger downregulation of mitochondrial-related pathways. (F) Expression profile plots of genes (mapped to human orthologue) related to glycolysis *(PFKP, PFKFB3, LDHB*, and *SLC16A3)*, thioredoxin (*TXN*), and the non-oxPPP (*TALDO1*). Gene expression was represented as z-scores across timepoint for the murine dataset (top) and the zebrafish dataset (bottom). Overall, there was a stronger concordance in expression patterns between zebrafish and KO mice compared to zebrafish and WT mice. Abbreviations: dpi (day post injury), GO (gene ontology), na (naive condition).

Given the technical differences between datasets, we hypothesized that conservation might be stronger at the pathway level rather than for individual genes. To test this, we compared GSEA across species, including wild-type murine RGCs (WT) at 2dpi, which do not regenerate, to distinguish injury-induced pathways in WT mice from those specifically driving regrowth in KO mice. All species/genotypes showed a significant upregulation of ribosome-related pathways and a downregulation of synapse-related pathways after injury (Fig. 5D-E). KO mice and zebrafish demonstrated similar upregulation of regeneration-related pathways, including IL-1 signaling, inflammation, PTEN stability, and macromolecule biosynthesis. Furthermore, both regenerative models showed an upregulation of the ER-Phagosome pathway and downregulation of dendrite spine development. A stronger downregulation of OXPHOS was found in zebrafish, while both mouse genotypes showed a higher downregulation of pathways related to cholesterol biosynthesis. Surprisingly, from broad GSEA analysis, only zebrafish showed a significant upregulation of glycolysis.

Since the rate of metabolic pathways is primarily regulated by rate-limiting steps, and the core enzymes of energy and biosynthetic metabolism are extremely highly conserved across all animal clades (Peregrín-Alvarez *et al*, 2009), we next compared the expression profile of the key metabolic enzymes identified above in zebrafish across all timepoints in the three models (Fig. 3; Fig. 5F). For glycolysis, we observed a comparable gene expression profile between zebrafish and KO mice (Fig. S2A), with upregulation of *PFKP* and *SLC16A3* early post-injury (Fig. 5F). *PFKFB3* and *LDHB* showed less pronounced expression changes in KO mice compared to zebrafish, but respectively a higher and lower expression as compared to WT mice (Fig. 5F). Similar agreements between zebrafish and KO mice were found for thioredoxin (TXN) and the non-oxPPP gene *TALDO1* (Fig. 5F), along with the upstream transcription factor *NFE2L2* (Fig. S2B). Zebrafish showed a stronger downregulation of genes related to OXPHOS as compared to both mouse genotypes (Fig. S2C). Regarding cholesterol-related genes, contrary to zebrafish, both WT and KO mice show a strong downregulation post-injury, but KO mice recovered expression at 21dpi, during axon elongation (Fig. S2D). In summary, zebrafish and KO RGCs displayed a comparable expression signature post-ONC, characterized by genes associated with enhanced glycolysis and thioredoxin-mediated detoxification. Therefore, these two pathways were selected as candidate evolutionarily conserved targets for validation.

### Inhibiting the thioredoxin antioxidant system or glycolysis delays axonal regeneration in adult zebrafish

To evaluate whether inhibiting thioredoxin detoxification interferes with RGC axonal regeneration, we blocked the entire system by using three inhibitors targeting its major components: 1) Conoidin A for Prdx (peroxiredoxin) (Haraldsen *et al*, 2009), 2) PX-12 for Txn (thioredoxin) (Ramanathan *et al*, 2007), and 3) auranofin for Txnrd (thioredoxin reductase) (Bindoli *et al*, 2009) (Fig. 6A). Since the sequencing data showed thioredoxin expression rising from 1dpi and peaking at 3dpi, we administered the inhibitor mix intravitreally at -1dpi, 1dpi, and 3dpi to cover the whole expression window (Fig. 6B). At 6dpi, axonal regeneration was quantified via biocytin anterograde tracing by measuring the fraction of reinnervated contralateral optic tectum (Beckers *et al*, 2019) (Fig. 6C). The 6dpi timepoint was selected as it corresponds to approximately 50% tectal reinnervation, making it ideal for detecting delays in axonal regrowth. A significant decrease in tectal reinnervation was observed in inhibitors-treated fish, which showed approximately 25% reinnervation (24.18 ± 2.03 SEM) compared to 60% in injured vehicle-treated fish (60.03 ± 2.18 SEM) (Fig. 6D). Of note, inhibitor treatment did not affect RGC survival in the retina (Fig. 6E; Fig. S3). Together, these data indicate that the thioredoxin system plays a critical role during axonal regeneration.

**Fig. 6.**
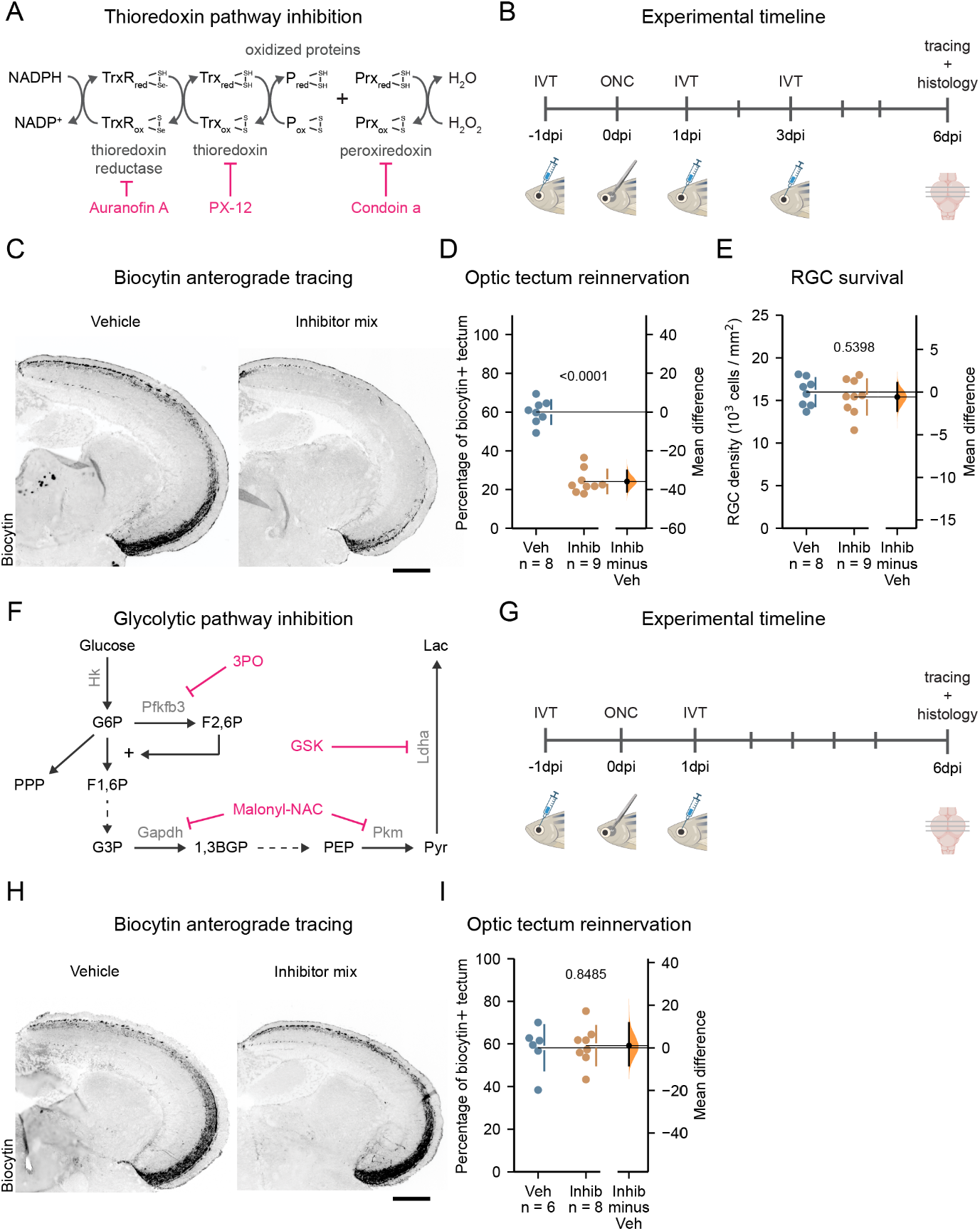
Inhibiting the thioredoxin antioxidant system significantly delays optic tectum reinnervation, while glycolysis inhibition does not. (A) Schematic overview of the thioredoxin antioxidant system and the inhibitors targeting different components of the system. (B) Timeline of the thioredoxin system inhibition experiment. Vehicle and inhibitor mix were injected at -1, 1, and 3dpi, and tectal reinnervation was evaluated at 6dpi. (C) Representative images of tectal reinnervation with thioredoxin system inhibition revealing reduced tectal reinnervation in Inhibitor-treated fish (right) at 6dpi. (D) At 6dpi, quantification of tectal reinnervation after thioredoxin system inhibition showed 60% reinnervation in vehicle-treated fish, compared to 25% in the inhibitor-treated group. (E) Quantification of RGC density revealed no significant reduction in RGC density with thioredoxin system inhibition at 6dpi. (F) Schematic overview of glycolysis and of the glycolytic enzymes targeted by the inhibitors. (G) Timeline of the glycolysis inhibition. Vehicle and inhibitors mix were injected at -1 and 1dpi, and tectal reinnervation was evaluated at 6dpi. (H) Representative images of tectal reinnervation with glycolysis inhibition show no delay in reinnervation at 6dpi compared to the control group. (I) Quantification of tectal reinnervation at 6dpi showed approximately 60% reinnervation in both the vehicle and inhibitor-treated groups. Data from n=8-9 (D-E) or n=6-8 (I) fish across two independent experiments, presented as mean ±SD and bootstrap 95% confidence interval versus vehicle-treaded fish. Scale bar 200 µm (C, H). Student’s T-test. P values are reported within the graphs. Abbreviations: IVT (interval injection), dpi (days post-injury).

Similarly, to inhibit glycolysis during the early regeneration phase, we used three inhibitors targeting the pathway at different levels: 3PO for Pfkfb3 (Emini Veseli *et al*, 2020), Malonyl-NAC for Gapdh and Pkm (glyceraldehyde-3-phosphate dehydrogenase) and Pkm (pyruvate kinase M) (Kulkarni *et al*, 2017), and GSK2837808A (GSK) for Ldha (lactate dehydrogenase A), to counteract the reduced expression of the opposing Ldhb (Billiard *et al*, 2013; Iwata *et al*, 2023) (Fig. 6F). Since sequencing data showed glycolysis upregulated until 3dpi, we administered inhibitors intravitreally at -1dpi and 1dpi (Fig. 6G). Tectal reinnervation was again evaluated at 6dpi. Contrary to thioredoxin inhibition, blocking glycolysis in the retina did not delay tectal reinnervation (Fig. 6H, I). Recent studies on *Pten* and *Socs3* co-deletion murine RGCs showed that, following axotomy in vitro, glycolysis is upregulated locally within regenerating axons but not in the soma (Masin *et al*, 2024). The lack of effect observed by inhibiting glycolysis via intravitreal injection might therefore be due to inability of the drugs to diffuse towards the outgrowing axons in the optic nerve. As pharmacological inhibition in the optic nerve is challenging, we opted for adult zebrafish retinal explants, cultured ex vivo.

We exposed the retinal explants to glycolysis-inhibiting media from the moment of seeding, thus allowing all cellular compartments of regenerating RGCs to be exposed to the inhibitor (Fig. 7A). Three conditions were adopted to comprehensively inhibit glycolysis: 2-DG (2-deoxy-D-glucose, 25 mM, a competitive inhibitor of glucose metabolism), galactose (25 mM, an alternative substrate to glucose), and the inhibitor mix used in vivo (3PO, Malonyl-NAC, and GSK, 9µM each) (Fig. 7B). 2-DG cannot be metabolized by the cell past the first step of glycolysis leading to accumulation and nearly complete inhibition of hexokinase (Hk) (Pajak *et al*, 2019). Galactose, when used instead of glucose, limits glycolysis, as it must be converted into glucose-6-phosphate (G6P) via the slow Leloir pathway before being further metabolized (Li *et al*, 2023; Robinson *et al*, 1992; Frey, 1996).

**Fig. 7.**
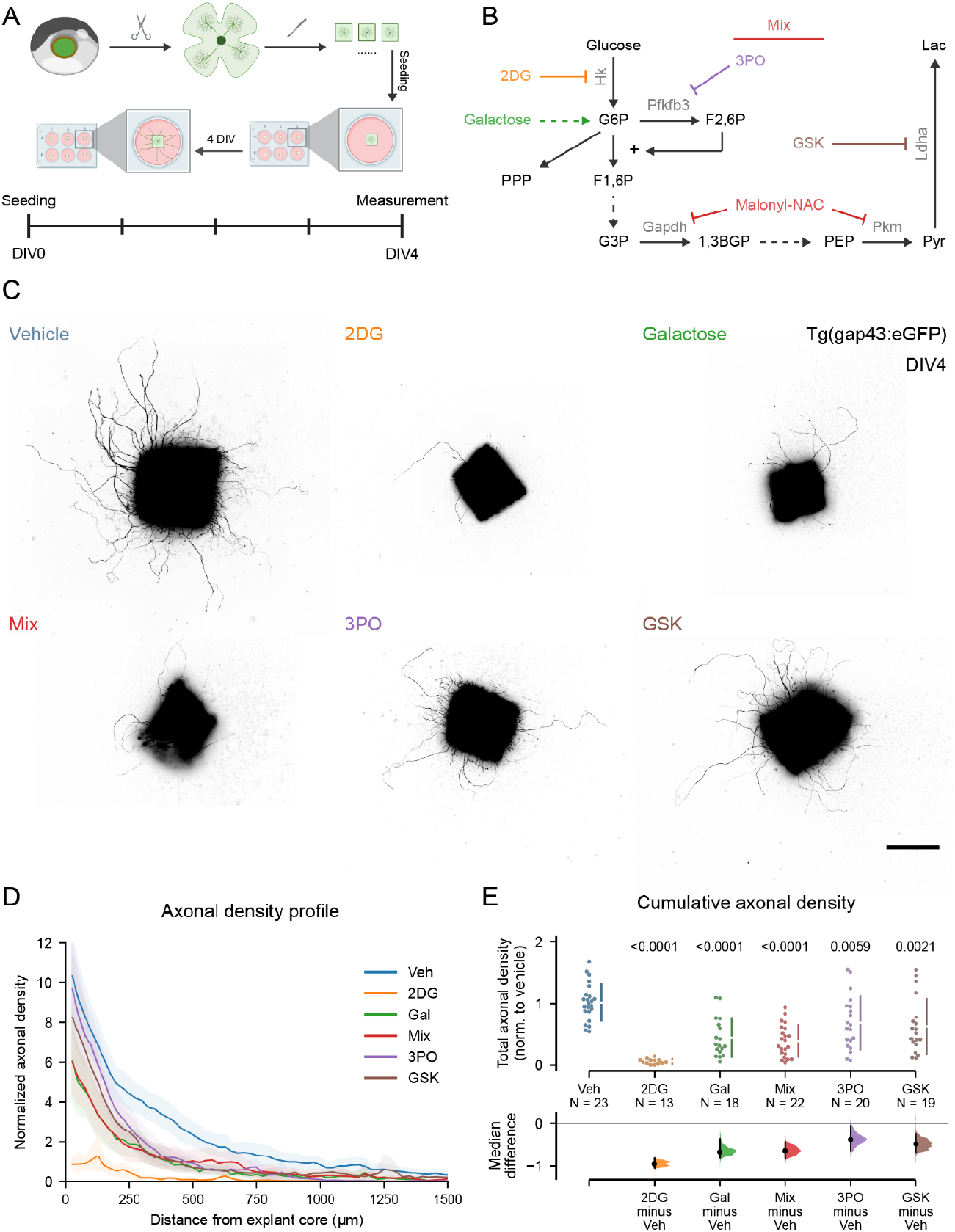
Inhibiting glycolysis significantly reduces axonal regeneration in adult zebrafish retinal explants. (A) Schematic overview of the experimental procedure. Retinas from naive *Tg(Tru*.*gap43:eGFP)*^*mil1*^ adult zebrafish were dissected, cut into explants and cultured, as described previously(Van houcke *et al*, 2017). After four days, regenerated axons were quantified. (B) Five glycolysis inhibition conditions were tested: (1) 2-DG (2-deoxy-D-glucose, orange); (2) Galactose (Gal, green); (3) an inhibitor mix (red, 9µM of 3PO, Malonyl-NAC, and GSK2837808A (GSK)); (4) 3PO alone (purple); (5) GSK alone (brown). (C) Representative images show that 2-DG caused the most severe axonal outgrowth reduction, with extremely few axons, growing only short distances. Galactose and inhibitor mix treatments also significantly impaired axonal outgrowth, but less severely. 3PO or GSK treatment induced a moderate reduction in axonal outgrowth compared to vehicle. Scale bar 200µm. (D) Cumulative distribution of axonal density profiles over distance from the explant. The 2DG treatment severely impaired both the density and length of axons. Galactose or inhibitor mix treatment caused a reduction in both sprouting and elongation, but less severe than 2-DG. Treatments with 3PO or GSK induced slightly reduced sprouting, but strongly impaired elongation. (E) Quantification of cumulative axonal density showed a 90% reduction with 2-DG treatment, around 60% reduction with galactose or inhibitor mix treatment, and approximately 40% reduction with 3PO or GSK treatment compared to the vehicle control. Data from N=13-23 explants, n=17 fish, across five independent experiments, presented as median ±95% confidence interval (D) or median ±SEM and bootstrap 95% confidence interval versus vehicle-treaded fish (E). One-way Kruskal-Wallis ANOVA (E). P values are reported within graphs. Abbreviations: DIV (day in vitro), Veh (vehicle). Figure containing assets from BioRender.com.

Unlike the inhibitor mix, 2-DG and galactose affect glucose metabolism upstream of the G6P step, where the PPP branches out from glycolysis (Fig. 3F), and therefore the PPP was also affected under these two conditions. Additionally, we individually inhibited Pfkfb3 with 3PO (9µM) and blocked Ldha with GSK (9µM). Axonal outgrowth of retinal explants was quantified at day in vitro 4 (DIV4). We found that all five treatments significantly affected both the axonal length (Fig. 7C-D) as well as the total axonal density (Fig. 7C, E), albeit with different magnitudes. 2-DG had the most severe effect, nearly abolishing neurite outgrowth (Fig. 7C-D). Treatment with galactose or the inhibitor mix resulted in a comparable reduction of approximately 60% in axonal outgrowth, and the regenerating axons were notably shorter than those in the vehicle group (Fig. 7C-D). 3PO or GSK treatment caused a more limited effect, but nonetheless significant, on both length and density of regenerating axons (Fig. 7C-D). Together, these data highlight that glycolysis plays an important role in axonal regeneration, and that its inhibition at all levels impairs regrowth. Complete blockade upstream of both glycolysis and the PPP causes a collapse of the outgrowth capacity in adult zebrafish, while specific targeting of glycolysis, both upstream and downstream, significantly impairs regeneration.

## Discussion

In this study, we identified that spontaneous axonal regeneration of RGCs upon ONC in adult zebrafish is accompanied by a profound alteration of their transcriptomic profile. In the early phases, during initiation of axonal regrowth and extension, we found a deep rearrangement of the expression of genes related to cellular energy metabolism, with OXHPOS downregulated and glycolysis and the pentose phosphate pathway (PPP) upregulated. This is accompanied by a specific upregulation of the thioredoxin system of antioxidant response. These expression signatures closely mirror those observed in the *Pten* and *Socs3* co-deletion murine model of induced axonal regeneration, suggesting these are evolutionarily conserved mechanism and potentially druggable targets. Inhibition of the thioredoxin system in vivo in adult zebrafish, via intravitreal injection, significantly disrupted tectal reinnervation, whereas inhibition of glycolysis did not. Conversely, inhibition of glycolysis in retinal explants cultured ex vivo, where axons were directly exposed to the drugs, significantly reduced axonal outgrowth, highlighting the critical role of glycolysis as a local axonal mechanism.

### The axonal regrowth process is defined by distinct gene expression phases

Analyzing the expression profile of RGCs during regrowth after ONC, we have found distinct phases of regeneration defined by characteristic transcriptomic signatures. Early on, at 1dpi, we found features characteristic of synaptic and dendritic pruning, aligning with the evidence that axonal regeneration of RGCs in adult zebrafish is preceded by a thinning of the IPL layer in the retina and loss of synaptic markers (Beckers *et al*, 2019). In this early phase, we also observed an immediate activation of regeneration-associated pathways, including the inflammation-related signaling of cytokines, such as IL-1. Later, the phase of axonal growth through the optic nerve (from 1 to 6dpi) is dominated by known signatures related to increased RNA biogenesis and translation. This is a central pathway regulated by the PIP3-AKT-mTOR axis (Beckers *et al*, 2019; Szwed *et al*, 2021). More interestingly, the phase of growth past the midline and of early tectal innervation (from 3 to 6dpi) is characterized by a unique signature of lipid biosynthesis (discussed below). Finally, the later phases of pre- and post-synaptic establishment and refinement return to a signature very close to that of non-injured neurons, with a downregulation of most regeneration-associated pathways.

### Regeneration is initiated by a profound reprogramming of energy metabolism

The regrowth of axons after injury is closely tied to energy production, and evidence shows that mitochondria are critical players in this. Their transport to the growth cone to fuel regeneration is observed in animal models and tissues capable of spontaneous regeneration (Han *et al*, 2016; Xu *et al*, 2017; Mar *et al*, 2014), and its activation in mammalian CNS models has been proven to induce axonal regrowth (Cartoni *et al*, 2017; Zhou *et al*, 2016). We previously demonstrated that dendritic pruning preceding axonal regrowth after ONC in adult zebrafish is accompanied by a relocation of mitochondria from the dendritic to the axonal compartment (Beckers *et al*, 2023b). Supporting this, here we found an upregulated expression of the Trak adaptor proteins, which link mitochondria to kinesin and dynein motors, allowing their transport along microtubules (van Spronsen *et al*, 2013). Interestingly, we found an upregulation of glycolytic genes early after injury and a pronounced downregulation of genes associated with OXPHOS throughout the regrowth process. The latter may represent a mechanism to mitigate oxidative stress caused by damaged axonal mitochondria (Zorov *et al*, 2014; Hill *et al*, 2016), with glycolysis compensating for the resulting energy deficit.

Recently, local upregulation of glycolysis within the axons was found to a key mechanism underlying the induced regrowth capability of *Pten* and *Socs3* co-deleted mouse RGCs (Sun *et al*, 2011; Masin *et al*, 2024). When comparing our transcriptomic data of adult zebrafish with those of *Pten* and *Socs3* co-deleted RGCs after ONC (Jacobi *et al*, 2022), we found that the overexpression of crucial rate-limiting enzymes of glycolytic genes is extremely conserved between the two models. Among others, phosphofructokinase 2 (PFK2, isoform PFKFB3), lactate dehydrogenase (LDH), and monocarboxylate transporter 4 (MCT4) particularly emerged as potential evolutionarily conserved targets. The latter two respectively mediate the reduction of pyruvate into lactate and its secretion from the cell. The generation of lactate is coupled with the oxidation of NADH to NAD^+^, an essential process for sustaining glycolysis when OXPHOS activity is restricted. Moreover, maintenance of NAD^+^ levels is known to be required to prevent degeneration of the axon (Conforti *et al*, 2014; Loreto *et al*, 2015). PFK2 acts in the first stages of glycolysis, increasing the activity of the main rate-limiting enzyme of the pathway, phosphofructokinase (PFK). In this study, the inhibition of glycolysis in vivo, via intravitreal injection, failed to affect the tectal reinnervation. Conversely, inhibiting glycolysis in the medium of explants cultured ex vivo, showed a severe impact on axonal outgrowth. Considering this, we postulate that enhanced glycolysis is a critical, evolutionarily conserved mechanism that is central to axonal regrowth. This notion is reinforced by evidence showing that also during development, glycolytic enzymes accumulate in the axon during outgrowth and that local inhibition of glycolysis causes growth cone collapse (Ketschek *et al*, 2021).

### Energy metabolism is coupled with macromolecule biosynthesis and antioxidant response

Glycolysis and mitochondria not only provide the cells with energy but are also critical in providing intermediates and cofactors that are crucial for many cellular processes. An important example is the PPP, which branches off from glycolysis to reduce NADPH and generate ribose sugars for nucleotide synthesis. In cells not requiring DNA synthesis for mitosis, the end products of the PPP are recycled back into glycolysis intermediates via the non-oxidative branch of the PPP. During the phase of axonal elongation, we found an upregulation of the enzymes transaldolase (TALDO) and transketolase (TK), which feed intermediates back into glycolysis. As the PPP subtracts intermediates that would otherwise be used for glycolysis, we hypothesize that this mechanism of upregulation of the non-ox PPP is aimed at allowing restorations of cellular NADPH via the oxPPP, while returning intermediates to glycolysis for energy production. This is known as “pentose overflow” and occurs in cells in which the requirement of NADPH exceeds the one of ribose, such as under oxidative stress (TeSlaa *et al*, 2023). Interestingly, we observed an upregulation of *tktb*, which encodes the isoform orthologue of the mammalian TKTL, the expression of which is characteristic of the metabolic adaptations of hypoxic cancerous cells (Baptista *et al*, 2022).

NADPH is a crucial cofactor for the biosynthesis of lipids and of polyprenylic chains by the mevalonate pathway (Chandel, 2021). These chains are used for prenylation of signaling proteins, such as the Ras and Rho family (Wang & Casey, 2016), as well as precursors for the synthesis of cholesterol, coenzyme Q and dolichols (Mullen *et al*, 2016). Cholesterol particularly plays an important role in the organization of lipid rafts on the neuronal membrane (Korade & Kenworthy, 2008; Cerqueira *et al*, 2016). Besides biosynthesis, NADPH also acts as an electron donor for the reductases of several antioxidant response systems, such as glutathione and thioredoxin (An *et al*, 2024). Among these, during axonal regeneration, we observed a strong upregulation of thioredoxin instead of others, such as superoxide dismutase or catalase. The thioredoxin system acts to reduce cysteine residues on proteins damaged by oxidation (An *et al*, 2024), but when coupled with peroxiredoxin, can also reduce H_2_O_2_ back to H_2_O (Rhee & Kil, 2017). It is unclear why the thioredoxin system becomes the most prevalent after injury and further research is needed to elucidate this. Inhibiting the entire system caused a significant decrease in optic tectum reinnervation at 6 dpi, but critically did not cause any detectable loss of RGCs in the retina, suggesting that this is a mechanism directly related to axonal regrowth rather than purely survival after axonal damage (Munemasa *et al*, 2008). In support of this hypothesis, thioredoxin was previously identified as a critical mediator of the enhanced axonal outgrowth observed after conditional lesion in *Xenopus* dorsal root ganglia neurons in vitro (Tonge *et al*, 2008). Thioredoxin is also found to be upregulated after ONC in regeneration-competent *Pten* and *Socs3* co-deletion RGCs and therefore is likely an evolutionarily conserved mechanism of axonal regeneration.

### Bridging the gap between fish and mammals: remaining issues and future perspectives

We found that early axonal regrowth in adult zebrafish shares metabolic features with the *Pten* and *Socs3* co-deletion murine model, including the upregulation of glycolysis and of the thioredoxin system. Zebrafish exhibited stronger downregulation of OXPHOS-related genes, although it remains unclear whether this plays a role in controlling ROS post-injury. Additionally, the mevalonate and cholesterol synthesis pathways were inversely regulated: upregulated in zebrafish during axonal elongation but rapidly downregulated in *Pten* and *Socs3* co-deleted RGCs after injury. Interestingly, expression of these pathways was recovered in co-deleted RGCs at 21 dpi, when axons cross the optic chiasm. Statin-mediated inhibition of these pathways is reported to promote axonal sprouting in murine CNS neurons, suggesting that suppression aids early sprouting, while activity may support later phases of regeneration (Tang, 2022; Li *et al*, 2016; Shabanzadeh *et al*, 2021).

Ample evidence shows that neuronal metabolic pathways can be regulated at the subcellular level (Wolfe *et al*, 2024; Masin *et al*, 2024; Ketschek *et al*, 2021), potentially driven in part by local translation within axons and dendrites (Glock *et al*, 2021; Shigeoka *et al*, 2016). In this study, we did not investigate whether the upregulated transcripts localize to specific compartments to promote localized phenotypes, and further research is needed to elucidate this. Nonetheless, based on our findings, we elect glycolysis and thioredoxin as key, conserved mechanisms of axonal regeneration in vertebrates. Targeting these pathways selectively, with precise temporal and subcellular delivery, could serve as a crucial steppingstone toward our ability to restore the capacity for regeneration in the adult mammalian CNS.

## Acknowledgments

We thank Evelien Herinckx and Arnold Van Den Eynde for the animal caretaking, Stephanie Mentens, Iene Kemps and Marijke Christiaens for technical support, and Julie De Schutter, Pieter-Jan Serneels for the critical discussions. S.B, A.V.D, and Lu.M. were supported by personal fellowships funded by the Research Foundation Flanders (FWO, Belgium) (fellowships 1165020N, 1S94218N, 1S42720N). A.B holds a personal L’Oréal/UNESCO (For Women in Science) fellowship. This research was funded by the FWO (project G082221N) and the KU Leuven Research Council (C14/22/074).

## Author contributions

Conceptualization: AZ, SB, LM, LuM; Data curation: AZ, LuM; Formal analysis: AZ, LuM; Funding acquisition: AZ, SB, AVD, AB, LM, LuM; Investigation: AZ; Methodology: AZ, SB, AVD, AB, LM, LuM; Project administration: AZ, LM, LuM; Resources: AZ, LM, LuM; Software: AZ, LuM; Validation: AZ, SB, LuM; Visualization: AZ, LM, LuM; Writing—original draft: AZ, LuM; Writing—review and editing: SB, LM.

## Resource availability

### Lead contact

Further information and requests for resources and reagents should be directed to and will be fulfilled by the lead contact, Lieve Moons

### Materials availability

All non-commercial reagents or fish lines used in this paper are available from the lead contact upon request.

### Data and code availability

- Raw and processed bulk RNA sequencing data are available for download through Gene Expression Omnibus (GEO) GSE289140: https://www.ncbi.nlm.nih.gov/geo/query/acc.cgi?acc=GSE289140.
- All original data reported in this paper will be shared by the lead contact upon request.
- This paper does not report original code.

Any additional information required to reanalyse the data reported in this paper is available from the lead contact upon request

## Methods and Methods

### Reagents and Tools Table

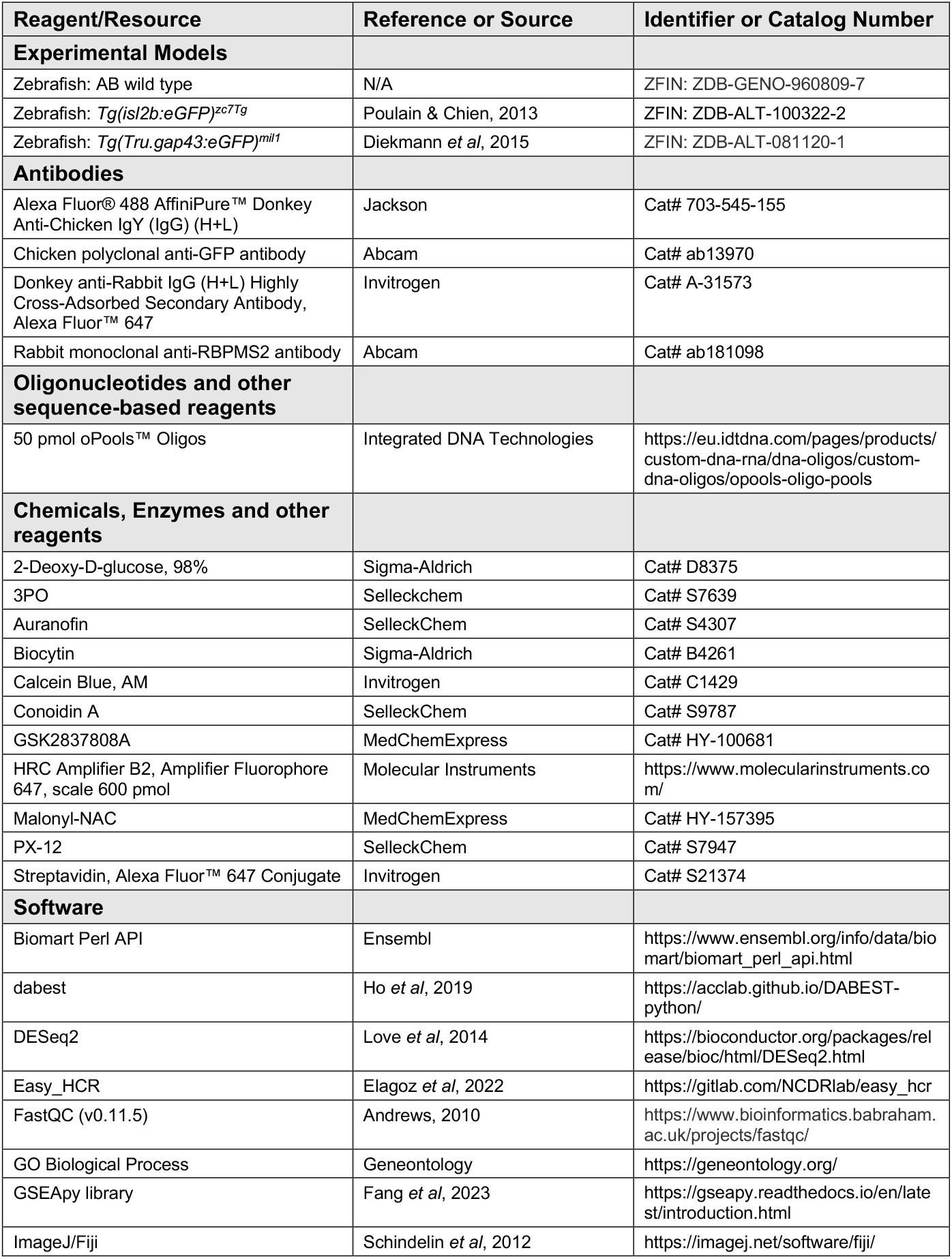
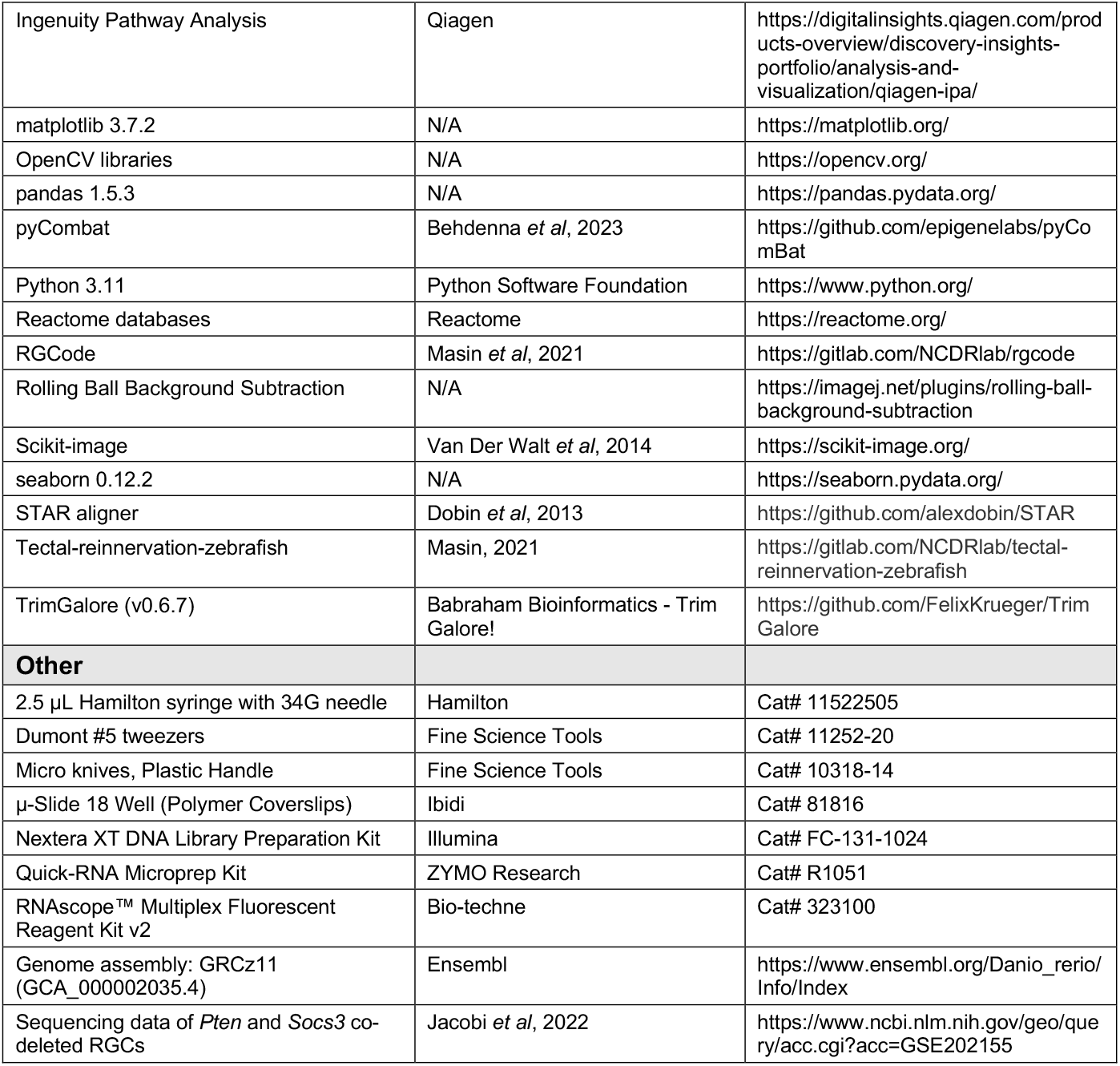

### Zebrafish maintenance

Zebrafish (*Danio rerio*) were housed in a standardized flow-through system with 15 fish per tank (ZebTec 3.5 L tanks) at 28°C in a 14 h light/10 h dark cycle. The fish were fed twice per day with dry food and brine shrimp. All experiments were performed on equally sized, 5-to 9-month-old adult zebrafish of both sexes. All the experiments were approved by KU Leuven Animal Ethics Committee (protocol number 111/2022; 226/2024) and executed in strict accordance with the European Communities Council Directive of 20 October 2010 (2010/63/EU). We used AB wild-type (WT), *Tg(isl2b:eGFP)*^*zc7Tg*^ (Poulain & Chien, 2013), in which the RGC-specific *isl2b* promoter drives the expression of eGFP, and *Tg(Tru*.*gap43:eGFP)*^*mil1*^, in which the *gap43* promoter drives the eGFP expression primarily in outgrowing neurons (Babraham Bioinformatics - Trim Galore!; Diekmann *et al*, 2015). In total, we used 31 AB WT fish, 123 *Tg(isl2b:eGFP)*^*zc7Tg*^, and 17 *Tg(Tru*.*gap43:eGFP)*^*mil1*^ fish for all experiments in this study. The exact number of fish used per experiment is depicted as individual data points in the figure graphs and/or mentioned in the corresponding figure legends.

### In vivo procedures

#### ONC injury

The optic nerve crush (ONC) injury was performed as previously described (Beckers *et al*, 2023a). Briefly, zebrafish were anesthetized with buffered 0.03% tricaine (MS-222, Millipore Sigma, Burlington, MA, USA) dissolved in system water. Connective tissue around the eye was carefully removed using fine forceps. Subsequently, the eye was gently elevated to expose the optic nerve. The optic nerve was then crushed for 10 seconds approximately 0.5 mm from the optic nerve head using Dumont #5 tweezers (Fine Science Tools, FST, Heidelberg, Germany). After the surgery, the eye was repositioned, and the fish were returned to the system water for recovery.

#### Thioredoxin antioxidant system inhibition in vivo

To study the effect of inhibiting the thioredoxin antioxidant system on zebrafish retinotectal regeneration post ONC injury, a combination of inhibitors targeting key enzymes of the thioredoxin pathway was injected intravitreally. The left eye of the fish was injected with 300 nL of 1 mM PX-12 (an inhibitor of thioredoxin), 1 mM auranofin (thioredoxin reductase inhibitor), and 0.05 mM conoidin A (a peroxiredoxin inhibitor) in dPBS (13.5% dimethylsulfoxide, DMSO) using a 2.5 μL Hamilton syringe with a 34G needle. To ensure the inhibition of the thioredoxin antioxidant system at the early regeneration stage, injections were conducted one day before ONC injury (-1dpi), one day and three days post ONC injury (1 and 3dpi). The vehicle control fish underwent injection and ONC at the same timepoint and were injected with the vehicle solution (13.5% DMSO in dPBS).

#### Glycolysis inhibition in vivo

Similarly, to evaluate the effect of inhibiting glycolysis on axonal regeneration of adult zebrafish RGCs post-ONC injury, 300 nL of an inhibitor mix containing 1 mM 3PO (Pfkfb3 inhibitor), 1 mM Malonyl-NAC (Gapdh and Pkm inhibitor), and GSK (Ldha inhibitor) in dPBS (13.5% DMSO) was injected intravitreally using a 2.5 μL Hamilton syringe with a 34G needle. To completely inhibit glycolysis during the early stages of regeneration, injections were performed 1 day before injury (-1dpi) and 1dpi. A 13.5% DMSO in dPBS solution was used as vehicle control.

#### Anterograde tracing of axonal regrowth and evaluation of RGC survival

Reinnervation of the optic tectum was evaluated using biocytin as an anterograde tracer, as previously described by Beckers et al.(2023a). At 5dpi, the optic nerve was transected between the optic nerve head and the crush site. A piece of gelfoam soaked in biocytin was then placed on the nerve stump. After three hours, the biocytin-traced eyes and brains were dissected. The biocytin in the brain was visualized on 10 μm-thick transverse cryosections by histochemical staining with streptavidin-Alexa 647 (1:300). The traced eyes were whole mounted, and stained for the Rbpms2b, an RGC marker (Kölsch *et al*, 2021). Briefly, the retinal whole mounts were permeabilized using a freeze-thaw step in 0.5% Triton X-100 in PBS (15 minutes, –80°C) and subsequently blocked with pre-immune donkey serum (PID, cat. no. S30-M, Sigma-Aldrich) for 2 hours at room temperature. The rabbit anti-RBPMS2 antibody (1: 200, 2% Triton X-100 in PBS with 10% PID) was incubated overnight at room temperature and visualized with an Alexa-conjugated secondary antibody (1:200, 2% Triton X-100 in PBS with 10% PID), which was incubated for 2 hours at room temperature.

### FACS sorting

*Tg(isl2b:eGFP)*^*zc7Tg*^ transgenic fish were used to sort the fluorescently labeled RGCs (Poulain & Chien, 2013; Kölsch *et al*, 2021). The single cell suspensions were obtained based on the protocol of Kölsch et al. (2021), with minor modification. Briefly, adult fish at naive condition and different timepoints post-injury (1, 3, 6, 10, and 14 days) were euthanized by submersion in buffered 0.1% tricaine (w/v in system water) (MS-222, MilliporeSigma, Burlington, MA, USA). The retinas of the fish were dissected in oxygenated Ames medium (cat. no. A1420, Sigma-Aldrich), and then incubated in 16 U/ml papain (cat. no. LS003118, Worthington) in oxygenated Ames together with 10 U/ml DNase (cat. no. LK003172, Worthington) for 45 min at 28°C. The digestion was stopped by rinsing the tissue with 1.5% ovomucoid (w/v, cat. no. LS003087, Worthington) in oxygenated Ames with 10 U/ml DNase, followed by another rinsing step with 0.4% bovine serum albumin (BSA, w/v, cat. no. A7906, Sigma-Aldrich) in oxygenated Ames with 10 U/ml DNase. The tissue was gently triturated with a 20-200 µL plastic pipette in the BSA solution. The obtained cell suspension was filtered through a 35 µm cell strainer. The AB wild-type fish were used to determine the background level of eGFP and adjust the gating. The cells were incubated with the live cell marker calcein blue (25 µM) to identify live cells. Living RGCs (defined as calcein blue + and eGFP+ cells) were isolated with fluorescence-activated cell sorting (FACS, Sony SH8000, Tokyo, Japan, 100 µm nozzle size, semi-purity sort mode, sample pressure 4). RGCs were collected in low adhesion Eppendorf tubes containing RNA lysis buffer (Zymo Research, Irvine, CA, USA) with RNAse inhibitor (80 U/mL, cat. no. 3335399001, Roche) and stored at ™80°C until RNA isolation. For each condition (naive and all timepoints post-injury), retinas from 123 adult fish were dissected. For each replicate, 10K RGCs were sequenced, with five to seven replicates obtained for per condition across 9 independent experiments.

### RNA Sequencing and bioinformatic analysis

#### RNA sequencing

RNA was extracted in two separate batches using the Quick-RNA Microprep kit according to the manufacturer’s instructions. The samples underwent bulk RNA sequencing in collaboration with the KU Leuven Genomics Core (https://www.genomicscore.be/). Libraries were prepared using the Smart-Seq2 method (RGC samples, Nextera XT DNA, Illumina, San Diego, CA, USA), and sequencing was performed on an Illumina HiSeq 4000 platform.

#### Alignment and quality control

Technical replicates of RNA-seq samples were merged post-sequencing, and adapter sequences were removed using TrimGalore (v0.6.7) (Babraham Bioinformatics - Trim Galore!). Quality control was conducted using FastQC (v0.11.5) (Andrews, 2010). Sequencing reads were aligned to the zebrafish reference genome (Ensembl-GRCz11) using the STAR aligner (Dobin *et al*, 2013). Genes with low expression were filtered out using a custom-developed R script, and batch effects were corrected with pyCombat (Behdenna *et al*, 2023).

#### Differential gene expression analysis

Differential gene expression analysis between conditions (naive condition vs. 1, 3, 6, 10, and 14dpi) was performed using DESeq2 (Love *et al*, 2014). Genes were defined as differentially expressed with minimally |log2FC| >1 and FDR <0.05. PCA plot was made using the DESeq2 package. To identify gene modules across the regeneration timeline, we collected all genes showing differential expression versus naive on at least one of the timepoint. For each, the normalized reads were z-scored across time to obtain expression profiles. The clustering of gene expression profiles was carried out using the scikit-learn library (Pedregosa FABIANPEDREGOSA *et al*, 2011). Briefly, the data was scaled, dimensionality was reduced using principal component analysis (PCA, 2 components) and finally clustering was performed via K-means clustering (k=5).

### Gene set enrichment and pathway analysis

To identify pathways enriched in different gene modules, we performed overrepresentation analysis via the GSEApy library (Fang *et al*, 2023), using the Enrichr API against the GO Biological Process database. A pathway with an adjusted p-value < 0.05 was considered significantly enriched. For pathway analysis at each timepoint, the differentially expressed genes were used to perform GSEA against the GO Biological Process and Reactome databases, using the DESeq2 “stat” metric as pre-ranking value. A pathway with a q-value < 0.05 was considered differentially regulated. Additionally, The Ingenuity Pathway Analysis tool (Qiagen, Redwood City, CA, USA) was employed to analyze the enrichment of molecular and functional gene networks among the differentially expressed genes (FDR < 0.01) at each post-injury.

#### Gene expression comparison against mouse

Transcriptomic data of wild-type (WT) and *Pten* and *Socs3* co-deleted RGCs (KO) were obtained from the Jacobi dataset (Jacobi *et al*, 2022), and previously re-analyzed as pseudo-bulk (Masin *et al*, 2024). To compare differentially expressed genes at 1dpi in zebrafish versus 2dpi in mouse, the gene list of both organisms was mapped to their respective human orthologue, using the Biomart API of GSEApy. Genes not mapping to any human gene were excluded from this analysis. Genes with an FDR < 0.05, |log2FC| > 1 and log2FC with the same sign were classified as “shared expression”, while genes with opposite sign were classified as “opposite expression”. Genes differentially expressed only in one of the two organisms, were classified as “mouse-only” and “zebrafish-only”, respectively. Pathway enrichment was performed on the genes with shared and opposite expression as described above. Pathway analysis in WT, KO mice and zebrafish was performed as described above, using the differentially expressed gene list mapped to human against the GO Biological Process and Reactome datasets. Finally, to compare gene expression profile, the z-scored, normalized reads from each organism were again mapped to their human orthologue and plotted across all timepoints.

### In situ hybridization chain reaction and RNAscope

The eyes from naive fish were collected alongside eyes from fish subjected to ONC injury at 1 and 3dpi and fixed in 4% paraformaldehyde (PFA) in PBS for one hour. Following three rinsing steps, the tissue was incubated in a sucrose series (10%, 20%, and 30% w/v in PBS) for at least 12 hours per incubation step. Subsequently, the eyes were embedded in 1.25% agarose (30% sucrose in PBS) and sectioned into 10 µm cryosections. Sections were collected on SuperFrost® Plus Slides (Fisher Scientific, cat. no. 12-550-15). Four biological replicates were included per condition.

The in situ hybridization chain reaction was performed based on HCR v3.0 as described by Choi et al. (2018), Elagoz et al. (2022), Bergmans et al. (2024). Briefly, probes were designed using Easy_HCR (https://gitlab.com/NCDRlab/easy_hcr) and purchased from Integrated DNA Technologies (IDT). Retinal cryosections were dried at 37°C and then fixed in 4% paraformaldehyde (PFA) for 3 hours. After washing with PBS treated with diethyl pyrocarbonate (DEPC), the tissue was permeabilized with 1% Tween-20. The sections were then treated with proteinase K (10 μg/mL, cat. no. 3115887001, Roche) for 10 minutes at 37°C, and the reaction was stopped by incubating with 4% PFA in PBS-DEPC for 10 minutes at room temperature. Following three rinses, sections were incubated in hybridization buffer containing 30% formamide, 0.75 M NaCl, 75 mM sodium citrate, 9 mM citric acid pH 6.0, 0.1% Tween20, 50 μg/mL heparin, 2% Denhardt’s solution, and 10% dextran for 30 minutes at 37°C. Probes were added at a final concentration of 6.67 nM in the probe hybridization buffer and allowed to hybridize overnight at 37°C. After rinsing three times, the slides were incubated in amplification buffer (0.75 M NaCl, 75 mM sodium citrate, 0.1% Tween20, 10% dextran) for 30 minutes at room temperature. Each hairpin (H1 and H2, HRC amplifiers) was separately prepared by heating 3 pmol at 95°C for 90 seconds and then cooling to room temperature over 30 minutes. Hairpins were then added to the slides at a concentration of 40 nM in amplification buffer and incubated overnight at room temperature. Nuclei were stained with DAPI, and the tissues were mounted with Mowiol®.

The RNAscope assay was performed using the RNAscope™ Multiplex Fluorescent Reagent Kit v2, in accordance with the manufacturer’s instructions. Briefly, the probes were designed and manufactured by Bio-Techne. Retinal cryosections were further fixed and dehydrated, following the fresh-frozen sample preparation and pretreatment workflow. After creating a hydrophobic barrier, the sections were treated with RNAscope Hydrogen Peroxide and Protease IV, all in accordance with the manufacturer’s instructions. The RNAscope Multiplex Fluorescent v2 Assay was then conducted with a dilution factor of 1:1500 for the TSA Vivid fluorophores. Following the development of the three channels of the HRP signal, the nuclei were stained with DAPI, and the tissues were mounted with Mowiol®.

### Glycolysis inhibition ex vivo

The adult zebrafish retinal explant culturing protocol was adapted from Van houcke *et al* (2017) and Van Dyck *et al* (2023). The ibidi µ-Slide 18 Well were used for explant cultures. Briefly, the wells were coated with Poly-L-Lysine solution (333 µg/mL, P6282, Sigma-Aldrich) and Laminin solution (1 mg/mL, cat. no. 23017015, Gibco) overnight in the fridge. Adult *Tg(Tru*.*gap43:eGFP)*^*mil1*^ fish were dark-adapted overnight to facilitate the removal of pigmented epithelial cells. Then fish were euthanized by submersion in buffered 0.1% tricaine (w/v in system water) (MS-222, MilliporeSigma, Burlington, MA, USA). The sclera, cornea, and lens were removed to expose the retina. The pigmented epithelium was removed by flushing with complete fish medium (CFM), composed of Leibovitz’s L-15 medium (cat. no. 11415064, Gibco), 2% fetal bovine serum (cat. no. 10270106, Gibco), 1% GlutaMAX (cat. no. 35050038, Gibco), 2% MEM essential amino acids (cat. no. 11130051, Gibco), 2% B-27 (v/v, cat. no. 17504044, Gibco), 25 mM D-glucose (cat. no. A2494001, Gibco), 25 mM HEPES (cat. no. 15630056, Gibco), 2% Penicillin/Streptomycin (v/v, cat. no. 15140122, Gibco), 0.05% amphotericin B (v/v, cat. no. 15290018, Gibco), 0.5% gentamicin (cat. no. G1272, Merck), adjusted pH to 7.4). The retina was further cut into approximately 0.1 mm^2^ pieces using micro knives. Nine explants were harvested per retina, and one explant was seeded per well with the RGC layer facing the bottom. The retinal explants from a single retina were distributed across wells with different conditions. For the DMSO control, the CFM medium was added with 0.03% DMSO. For the 2-Deoxy-D-glucose (2-DG) condition, the CFM was supplemented with 25 mM 2-DG (with 0.03% DMSO). For the Galactose condition, we used CFM without glucose and brought the galactose concentration in the medium to 25 mM (with 0.03% DMSO). For glycolysis inhibition, an inhibitor mix of 9µM 3PO (an inhibitor for Pfkfb3, cat. no. S7639, Selleckchem), 9µM Malonyl-NAC (an inhibitor for Gapdh and Pkm), and 9µM GSK (an inhibitor for Ldha, cat. no. HY-100681, MCE) were added to the CFM medium (with 0.03% DMSO).

At day one in culture, the medium was changed from CFM to complete Neurobasal-A medium (CNBA), composed of Neurobasal™-A Medium (no D-glucose, no sodium pyruvate, cat. no. A2477501, Gibco), 25mg/L sodium pyruvate (cat. no. 11360070, Gibco), 25mM Glucose (cat. no. A2494001, Gibco), 2% B-27 (v/v, cat. no. 17504044, Gibco), 1% L-glutamine (v/v, cat. no. 25030081, Gibco), 2.5 mM HEPES, 2% Penicillin/Streptomycin (v/v, cat. no. 15140122, Gibco), 0.05% amphotericin B (v/v, cat. no. 15290018, Gibco), 0.5% gentamicin (cat. no. G1272, Merck). For the vehicle control, the CNBA medium was modified to contain 0.03% DMSO. For the 2-DG condition, the CNBA medium was supplemented with 25 mM 2-DG (with 0.03% DMSO). For the Galactose condition, we used CNBA medium without glucose and added galactose to the medium at a concentration of 25 mM (with 0.03% DMSO). For glycolysis inhibition, 9µM 3PO (an inhibitor for PFKFB3) and 9µM GSK (an inhibitor for LDHA) were added to the CNBA medium (with 0.03% DMSO).

Half of the medium was refreshed daily. The explants were cultured in a humidified incubator with 5% CO_2_ at 28°C. On day 4 in vitro, the explants were fixed with 4% PFA in PBS for 30 minutes. After three rinses with PBS, the explants were permeabilized with 0.2% PBST (0.2% v/v triton X-100 in PBS) and then incubated with 20% PID (cat. no. S30-M, Sigma-Aldrich) in 0.2% PBST overnight. Following blocking, the explants were incubated with chicken anti-GFP (1:500) in 0.5% PBST (containing 10% PID) overnight and rinsed three times with 0.2% PBST. Next, the explants were incubated with donkey anti-chicken Alexa Fluor 488 (1:300) in 0.5% PBST (containing 10% PID) for 2 hours. After three rinses with PBST, nuclear staining was performed with DAPI (1:1000 in PBS), followed by a final rinse with PBS and mounting with Mowiol®.

### Imaging

#### HCR and RNAscope imaging

Retinal sections labeled with HCR or RNAscope were imaged using a Zeiss LSM900 Airyscan microscope equipped with a Plan-Apochromat 20× 0.8 NA objective in Airyscan CO-2Y mode, maintaining consistent imaging settings for the same batch of experiments. Images were acquired as mosaic Z-stacks covering the entire thickness of the section and visualized as maximum intensity projections.

#### Imaging of optic tectum reinnervation

Histological images of optic tectum sections were captured using a Leica DM6 microscope with a HC PL FLUOTAR L 20×/0.40 CORR objective, maintaining consistent imaging settings for the same batch of experiments.

#### Imaging of retinal whole mount for RGC counting

Immunohistochemical images of retinal whole mounts stained for Rbpms2b were captured using a Leica DM6 microscope with a HC PL FLUOTAR L 20×/0.40 CORR objective.

#### Imaging of retinal explants

Explant images were acquired using a Zeiss LSM900 in epifluorescence mode with a Plan-Apochromat 20×/0.8 M27 objective. They were captured as a mosaic to cover the entire field of regenerated axons.

### Image analysis

#### Quantification of optic tectum reinnervation

Quantification of tectal reinnervation was performed using a custom ImageJ script (Masin, 2021), as described by Beckers et al. (2019). Briefly, the area between the stratum opticum (SO) and the stratum fibrosum et griseum superficiale (SFGS) of the optic tectum was delineated, and the total area of the SFGS and SO was measured. The biocytin-positive area within the SFGS and SO was manually thresholded, and the biocytin-positive area was then measured. The percentage of reinnervation was defined as the biocytin-positive area divided by the total delineated area. For each fish, the reinnervation percentage was analyzed in at least three sections of the central optic tectum. The average reinnervation percentage for the sections from the same fish was calculated and presented as one data point, with six to nine animals used per experimental condition.

#### Quantification of RGC survival

The survival of RGCs was quantified using RGCode (Masin *et al*, 2021). Briefly, a model was trained on manual annotated images of Rbpms2 staining, and the number of RGCs was measured together with the area of the retina, to yield the density of RGCs.

#### Quantification of axonal outgrowth in retinal explants

The axonal outgrowth of retinal explants cultured ex vivo was quantified based on Gaublomme et al. (2013), with minor modifications. Image analysis was performed in ImageJ and in python using scikit-image (Van Der Walt *et al*, 2014) and OpenCV libraries (OpenCV). Briefly, the core of the explant was identified by thresholding the DAPI signal (Li algorithm). The axonal GFP signal was 1) background-subtracted (rolling ball algorithm), 2) smoothened with a Gaussian blur (σ=2), and 3) binarized with a Li thresholding. The “analyze particle” function of ImageJ was used to clean the binary image from impurities. Particles with a size > 1000 pixels and circularity > 0.1 were excluded from the binary imaging. The radius of the explant core was increasingly dilated in bins of 25 µm. Per each bin, the GFP+ axonal area was divided by the total area of the bin to obtain the axonal density. The profile of density across distance from the explant core was normalized by the mean density of the first three bins and by the size of the explant core. The cumulative density was obtained as the area under the curve of the density profile, either for the whole profile or in bins of 0-250 µm, 250-500 µm and > 500 µm of distance from the explant core. When accumulating data from independent experiments, the data of each experiment was normalized by the median of their relative vehicle control and expressed as fold-change.

### Statistical analysis

Detailed information on the number of samples and biological replicates used are reported in the legends of the figures. The number of n fish per condition/timepoint is reported in the graph under each group. For the ex vivo experiments, the number of N explants used for the experiment, from N fish, accumulated over several independent experiments are reported in the figure legends. Raw micrographs, acquired without saturation, were used for all data analyses. For visualization in figure panels, some images were inverted and/or contrast-enhanced by adjusting the white point or, when inverted, increasing the black point. The same degree of enhancement was consistently applied across all displayed images for comparative purposes. To display the axons of explants, the representative images were purposefully saturated to reliably show all axons. Statistical tests, such as ANOVA and t-tests, were performed alongside bootstrapping for significance assessment, with specific details provided in the figure legends. Estimation statistics with bootstrapping were employed due to their robustness with large sample sizes. The median was used in conjunction with Kruskal-Wallis ANOVA for datasets in which any group failed the Shapiro-Wilk normality test. Otherwise, the mean and Welch ANOVA were applied. All numerical data processing, visualization, statistical analyses, and bootstrapping were carried out in Python using the pandas, seaborn, matplotlib, scipy, and dabest (Ho *et al*, 2019) libraries. A p-value of <0.05 was considered statistically significant.

## Supplemental material

Table.S1 is related to Fig. 2E, containing all the genes in each module.

Table.S2 contains differentially expressed genes at all time points post-injury compared to the naive condition, including log2FoldChange, adjusted p-values, and statistical values for each time point.

Fig. S1, related to Fig. 1, shows the gating on the FACS density plot for isolating live RGCs for sequencing. Fig. S2, related to Fig. 5, shows the expression of key metabolic genes in zebrafish, regeneration-competent *Pten* and *Socs3* co-deleted mice, and regeneration-incompetent wild-type mice after optic nerve crush injury. Fig. S3, related to Fig. 6, shows representative images of RBPMS2 staining to identify zebrafish RGCs and their density after injury in vehicle- and inhibitor-treated retinas. Fig. S4, related to Fig. 7, reports the cumulative axonal density in short-, mid-, and long-range distances from the explant core.

**Fig S1.**
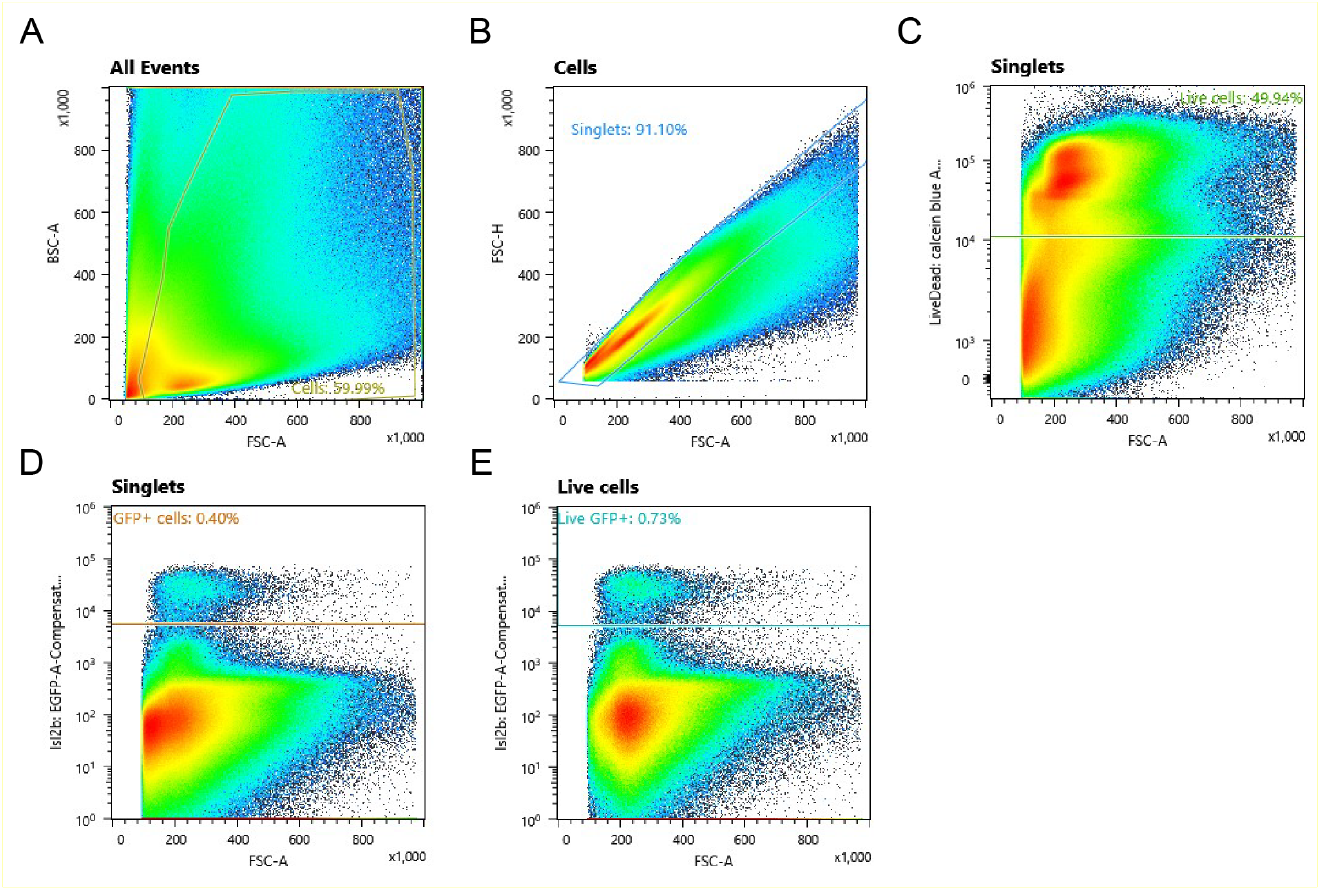
FACS gating strategy for sorting live adult zebrafish RGCs using Tg(isl2b:eGFP)^zc7Tg^. (A) Density plot showing all detected events based on forward scatter area (FSC-A) and back scatter area (BSC-A). The population with larger FSC-A values and higher complexity (BSC-A) was selected as cells to exclude debris and non-cellular particles. (B) Singlets were gated by selecting the cell population with a linear correlation between forward scatter area (FSC-A) and forward scatter height (FSC-H) to eliminate doublets and cell aggregates. (C) Live cells were identified from singlets using calcein blue staining (live cell marker). The calcein blue positive gate was set based on a negative control (unstained cell suspension), ensuring that only viable cells were selected. (D) RGCs were identified from singlets based on the endogenous GFP expression, as RGCs are the only retinal cell type expressing *isl2b*. The GFP positive gate was set based on a negative control (wild-type fish). (E) RGCs expressing GFP were selected from the live singlet population. The GFP positive gate was set based on a negative control (wild-type fish). The calcein blue and GFP double-positive RGCs were collected for further sequencing. Abbreviations: FSC-A (Forward Scatter-Area), BSC-A (Back Scatter-Area), FSC-H (Forward Scatter-Height).

**Fig S2.**
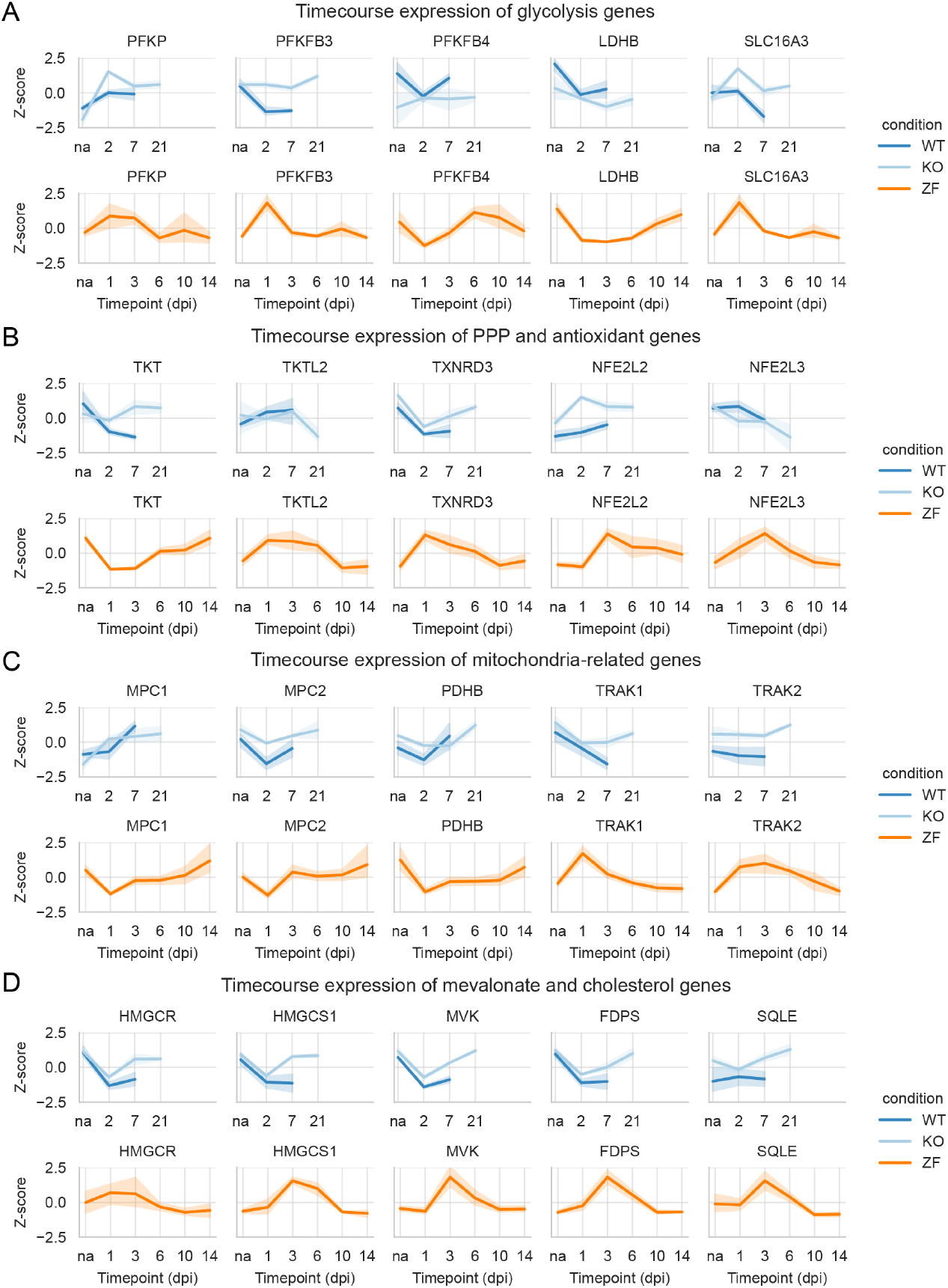
The overexpression of glycolysis and the thioredoxin system is conserved across regeneration-competent vertebrate models, while expression of OXPHOS and cholesterol biosynthesis are not. (A) Expression profile plots of genes (mapped to their human orthologue) related to glycolysis. Gene expression was represented as z-scores across timepoint for the murine dataset (top) and the zebrafish dataset (bottom). Overall, expression patterns showed greater similarity between zebrafish and *Pten* and *Socs3* co-deleted mice (KO) compared to zebrafish and wild-type (WT) mice. (B) Expression profile plots of genes (mapped to their human orthologue) related to the non-oxPPP (*TKT, TKTL2*), thioredoxin (*TXNRD3*) and antioxidant transcription factors (*NFE2L2, NFE2L3*). Gene expression was represented as z-scores across timepoint for the murine dataset (top) and the zebrafish dataset (bottom). Genes of the PPP and NFE2L2 showed a strong agreement in expression patterns between zebrafish and KO mice, while *TXNRD3* and *NFE2L3* did not. (C) Expression profile plots of genes (mapped to their human orthologue) related to the import of pyruvate into the mitochondrion and the Krebs cycle (*MPC1-2* and *PDHB*) and related to mitochondrial transport (*TRAK1-2*). Gene expression was represented as z-scores across timepoint for the murine dataset (top) and the zebrafish dataset (bottom). Agreement in the expression between zebrafish and mice was poor, with the exception of *MPC2* and, to a lesser extent, *PDHB*, suggesting that the conservation of genes related to mitochondria was more limited. (D) Expression profile plots of genes (mapped to their human orthologue) related to the synthesis of mevalonate and cholesterol. Gene expression was represented as z-scores across timepoint for the murine dataset (top) and the zebrafish dataset (bottom). Zebrafish and mice showed an opposite response, with the first upregulating expression during axon elongation, and mice downregulating it after injury. Nonetheless, expression was recovered in later phases of regeneration in KO mice. Abbreviations: dpi (day post injury), non-oxPPP (non-oxidative pentose phosphate pathway), na (naive), KO (*Pten*^*-/-*^; *Socs3*^*-/-*^, *Pten* and *Socs3* co-deleted), wild-type (WT), zebrafish (ZF).

**Fig S3.**
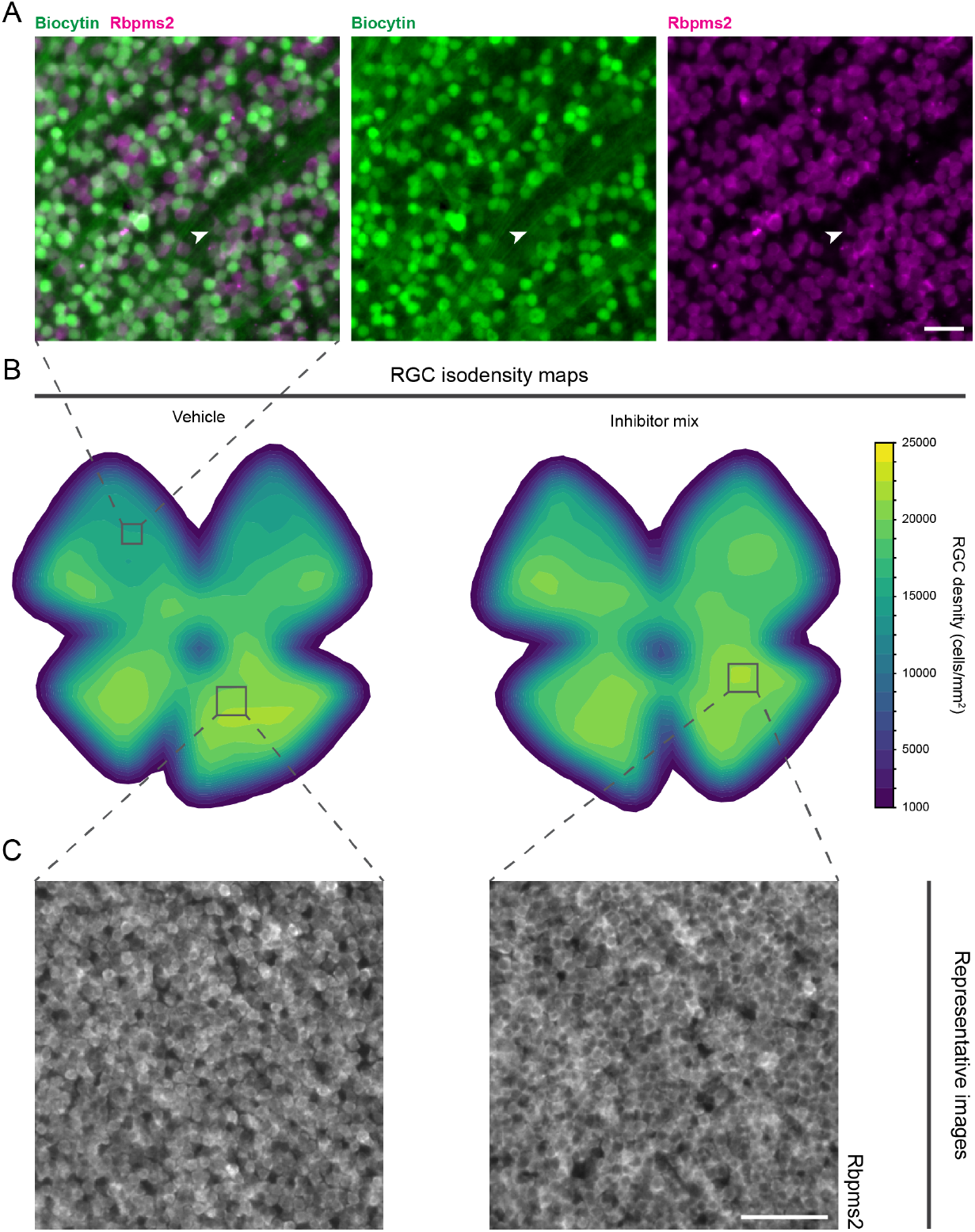
Inhibition of the thioredoxin system does not affect RGC survival after optic nerve crush. (A) Representative images from uninjured wholemount retinas immunostained for biocytin (retrograde tracing of RGCs, green) and Rbpms2 (magenta). The latter results in a more homogenous labelling of RGCs that is not obstructed by traced axons (arrowheads) and therefore constitutes a better option to reliably quantify all RGCs. Scale bar 20 µm. (B) Isodensity maps of Rbpms2-stained wholemount retinas for the quantification of RGC survival after ONC and vehicle or thioredoxin inhibition treatment. Vehicle or the inhibitor mix was injected at -1, 1, and 3dpi, and RGC survival was evaluated at 6dpi. No appreciable reduction in RGC density was observed upon inhibitor administration. (C) Representative images of the Rbpms2-immunostained retinas at 6dpi for the quantification of RGC survival after ONC and vehicle or thioredoxin inhibition treatment. No appreciable reduction in RGC density was observed upon inhibitor administration. Scale bar 50µm.

**Fig S4.**
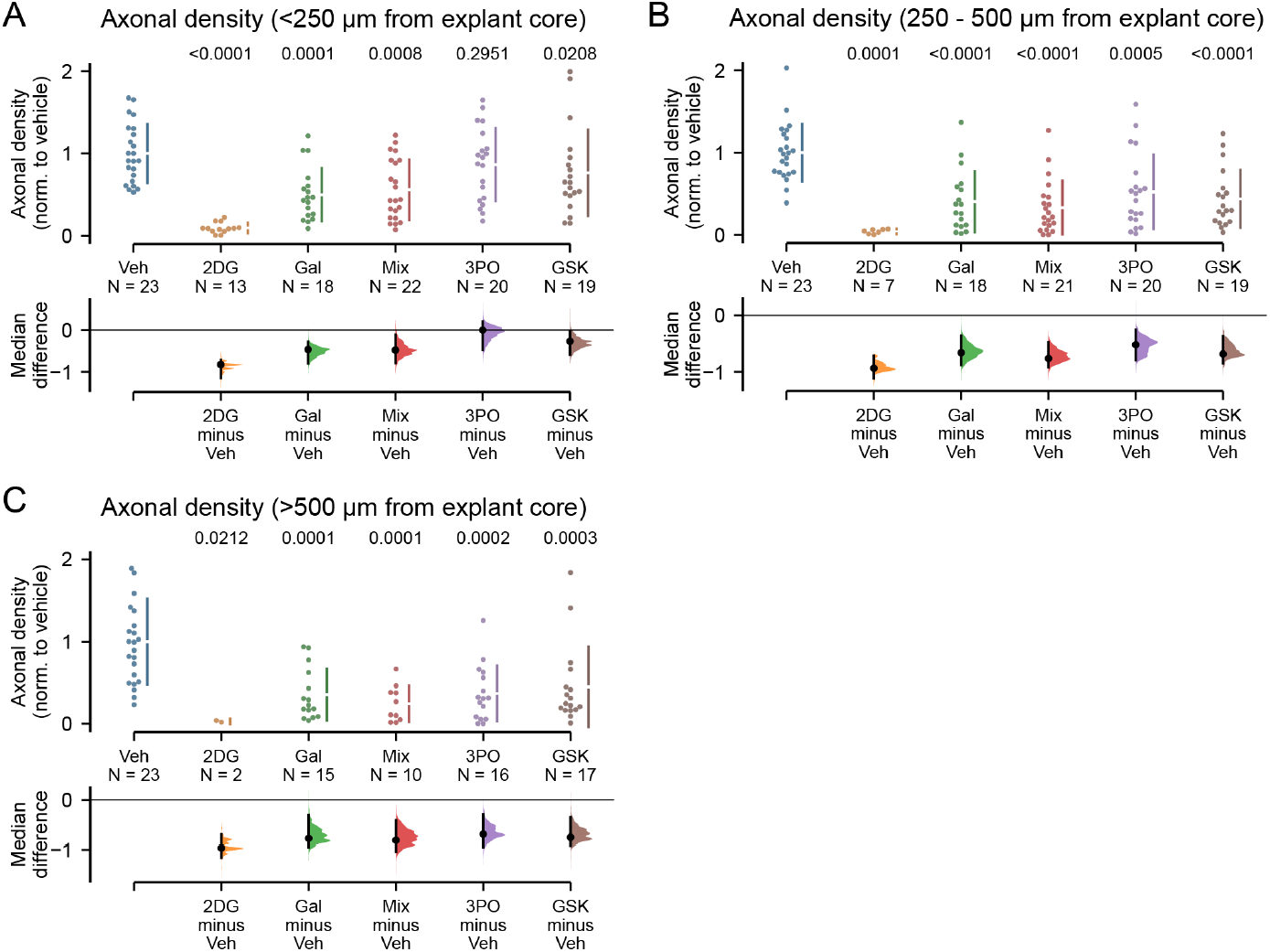
Inhibiting glycolysis significantly reduces long-range axonal regrowth in adult zebrafish retinal explants. (A) Retinas from adult *Tg(Tru*.*gap43:eGFP)*^*mil1*^ zebrafish were dissected and cut into explants, which were seeded in wells with the RGC layer facing down. Quantification of the cumulative axonal density within 250 µm from the explant core (short-range growth) revealed that 2-DG treatment resulted in an approximately 80% reduction in axonal density. Galactose and inhibitor mix treatments caused around 45% total axonal density reduction. Treatment with GSK exhibited a reduction of approximately 25% as compared to vehicle control. In contrast, 3PO did not significantly reduce axonal density. (B) Quantification of the cumulative axonal density in the range from 250 µm to 500 µm from the explant core (mid-range growth) showed that 2-DG treatment led to an approximately 95% reduction in axonal density. Galactose and inhibitor mix treatments resulted in roughly 65% and 75% reductions, respectively. Treatments with 3PO or GSK resulted in reductions of approximately 55% and 70%, respectively, as compared to the vehicle control. (C) Quantification of the cumulative axonal density in the range further than 500 µm from the explant core (long-range growth) revealed that 2-DG treatment resulted in an approximately 95% reduction in axonal density. Galactose and inhibitor mix treatments induced a reduction of around 80% total axonal density, while treatments with 3PO or GSK caused a reduction of approximately 70% as compared to the vehicle control. Data from N=13-23 explants, n=17 fish, across five independent experiments, presented as median ± SEM and bootstrap 95% confidence interval versus vehicle-treaded fish. One-way Kruskal-Wallis ANOVA. P values are reported within the graphs. Abbreviations: Veh (vehicle).

## References

Akram R, Anwar H, Javed MS, Rasul A, Imran A, Malik SA, Raza C, Khan IU, Sajid F, Iman T, et al (2022) Axonal regeneration: underlying molecular mechanisms and potential therapeutic targets. Biomedicines 10: 3186

An X, Yu W, Liu J, Tang D, Yang L, Chen X (2024) Oxidative cell death in cancer: mechanisms and therapeutic opportunities. Cell Death Dis 2024 15:8 15: 1–20

Andrews S (2010) FastQC: a quality control tool for high throughput sequence data. https://www.bioinformatics.babraham.ac.uk/projects/fastqc/

Andries L, Van Hove I, Moons L, De Groef L (2016) Matrix metalloproteinases during axonal regeneration, a multifactorial role from start to finish. Mol Neurobiol 2016 54:3 54: 2114–2125

Babraham Bioinformatics Trim Galore!. https://www.bioinformatics.babraham.ac.uk/projects/trim_galore/.

Baptista I, Karakitsou E, Cazier JB, Günther UL, Marin S, Cascante M (2022) TKTL1 knockdown impairs hypoxia-induced glucose-6-phosphate dehydrogenase and glyceraldehyde-3-phosphate dehydrogenase overexpression. Int J Mol Sci 23: 3574

Beckers A, Bergmans S, Van Dyck A, Moons L (2023a) Analysis of axonal regrowth and dendritic remodeling after optic nerve crush in adult zebrafish. Methods Mol Biol 2636: 163–190

Beckers A, Van Dyck A, Bollaerts I, Van houcke J, Lefevere E, Andries L, Agostinone J, Van Hove I, Di Polo A, Lemmens K, et al (2019) An antagonistic axon-dendrite interplay enables efficient neuronal repair in the adult zebrafish central nervous system. Mol Neurobiol 56: 3175–3192

Beckers A, Masin L, Van Dyck A, Bergmans S, Vanhunsel S, Zhang A, Verreet T, Poulain FE, Farrow K, Moons L (2023b) Optic nerve injury-induced regeneration in the adult zebrafish is accompanied by spatiotemporal changes in mitochondrial dynamics. Neural Regen Res 18: 219–225

Beckers A, Vanhunsel S, Van Dyck A, Bergmans S, Masin L, Moons L (2021) Injury-induced autophagy delays axonal regeneration after optic nerve damage in adult zebrafish. Neuroscience 470: 52–69

Behdenna A, Colange M, Haziza J, Gema A, Appé G, Azencott CA, Nordor A (2023) pyComBat, a Python tool for batch effects correction in high-throughput molecular data using empirical Bayes methods. BMC Bioinformatics 24: 1–9

Belin S, Nawabi H, Wang C, Tang S, Latremoliere A, Warren P, Schorle H, Uncu C, Woolf CJ, He Z, et al (2015) Injury-induced decline of intrinsic regenerative ability revealed by quantitative proteomics. Neuron 86: 1000

Bergmans S, Noel NCL, Masin L, Harding EG, Krzywańska AM, De Schutter JD, Ayana R, Hu CK, Arckens L, Ruzycki PA, et al (2024) Age-related dysregulation of the retinal transcriptome in African turquoise killifish. Aging Cell 23: e14192

Billiard J, Dennison JB, Briand J, Annan RS, Chai D, Colón M, Dodson CS, Gilbert SA, Greshock J, Jing J, et al (2013) Quinoline 3-sulfonamides inhibit lactate dehydrogenase A and reverse aerobic glycolysis in cancer cells. Cancer Metab 1: 1–17

Bindoli A, Rigobello MP, Scutari G, Gabbiani C, Casini A, Messori L (2009) Thioredoxin reductase: A target for gold compounds acting as potential anticancer drugs. Coord Chem Rev 253: 1692–1707

Bollaerts I, Veys L, Geeraerts E, Andries L, De Groef L, Buyens T, Salinas-Navarro M, Moons L, Van Hove I (2017) Complementary research models and methods to study axonal regeneration in the vertebrate retinofugal system. Brain Struct Funct 2017 223:2 223: 545–567

Cartoni R, Pekkurnaz G, Wang C, Schwarz TL, He Z (2017) A high mitochondrial transport rate characterizes CNS neurons with high axonal regeneration capacity. PLoS One 12

Cerqueira NMFSA, Oliveira EF, Gesto DS, Santos-Martins D, Moreira C, Moorthy HN, Ramos MJ, Fernandes PA (2016) Cholesterol biosynthesis: a mechanistic overview. Biochemistry 55: 5483–5506

Chandel NS (2021) Lipid metabolism. Cold Spring Harb Perspect Biol 13: a040576

Choi HMT, Schwarzkopf M, Fornace ME, Acharya A, Artavanis G, Stegmaier J, Cunha A, Pierce NA (2018) Third-generation in situ hybridization chain reaction: Multiplexed, quantitative, sensitive, versatile, robust. Development (Cambridge) 145

Conforti L, Gilley J, Coleman MP (2014) Wallerian degeneration: an emerging axon death pathway linking injury and disease. Nat Rev Neurosci 2014 15:6 15: 394–409

Dhara SP, Rau A, Flister MJ, Recka NM, Laiosa MD, Auer PL, Udvadia AJ (2019) Cellular reprogramming for successful CNS axon regeneration is driven by a temporally changing cast of transcription factors. Sci Rep 2019 9:1 9: 1–12

Diekmann H, Kalbhen P, Fischer D (2015) Characterization of optic nerve regeneration using transgenic zebrafish. Front Cell Neurosci 9: 130266

Divakaruni AS, Wallace M, Buren C, Martyniuk K, Andreyev AY, Li E, Fields JA, Cordes T, Reynolds IJ, Bloodgood BL, et al (2017) Inhibition of the mitochondrial pyruvate carrier protects from excitotoxic neuronal death. J Cell Biol 216: 1091–1105

Dobin A, Davis CA, Schlesinger F, Drenkow J, Zaleski C, Jha S, Batut P, Chaisson M, Gingeras TR (2013) STAR: ultrafast universal RNA-seq aligner. Bioinformatics 29: 15–21

Van Dyck A, Masin L, Bergmans S, Schevenels G, Beckers A, Vanhollebeke B, Moons L (2023) A new microfluidic model to study dendritic remodeling and mitochondrial dynamics during axonal regeneration of adult zebrafish retinal neurons. Front Mol Neurosci 16: 1196504

Elagoz AM, Styfhals R, Maccuro S, Masin L, Moons L, Seuntjens E (2022) Optimization of whole mount RNA multiplexed in situ hybridization chain reaction with immunohistochemistry, clearing and imaging to visualize octopus neurogenesis. bioRxiv: 2022.02.24.481749

Emini Veseli B, Perrotta P, Van Wielendaele P, Lambeir AM, Abdali A, Bellosta S, Monaco G, Bultynck G, Martinet W, De Meyer GRY (2020) Small molecule 3PO inhibits glycolysis but does not bind to 6-phosphofructo-2-kinase/fructose-2,6-bisphosphatase-3 (PFKFB3). FEBS Lett 594: 3067–3075

Fang Z, Liu X, Peltz G (2023) GSEApy: a comprehensive package for performing gene set enrichment analysis in Python. Bioinformatics 39

Fawcett JW (2020) The struggle to make CNS axons regenerate: why has it been so difficult? Neurochem Res 45: 144

Frey PA (1996) The Leloir pathway: a mechanistic imperative for three enzymes to change the stereochemical configuration of a single carbon in galactose. FASEB J 10: 461–470

Gagliardi D, Pagliari E, Meneri M, Melzi V, Rizzo F, Comi G Pietro, Corti S, Taiana M, Nizzardo M (2022) Stathmins and motor neuron diseases: pathophysiology and therapeutic targets. Biomedicines 10: 711

Gaublomme D, Buyens T, Moons L (2013) Automated analysis of neurite outgrowth in mouse retinal explants. J Biomol Screen 18: 534–543

Giacomello M, Pyakurel A, Glytsou C, Scorrano L (2020) The cell biology of mitochondrial membrane dynamics. Nat Rev Mol Cell Biol 2020 21:4 21: 204–224

Glock C, Biever A, Tushev G, Nassim-Assir B, Kao A, Bartnik I, Dieck S tom, Schuman EM (2021) The translatome of neuronal cell bodies, dendrites, and axons. Proc Natl Acad Sci U S A 118: e2113929118

Han Q, Xie Y, Ordaz JD, Huh AJ, Huang N, Wu W, Liu N, Chamberlain KA, Sheng ZH, Xu XM (2020) Restoring cellular energetics promotes axon regeneration and functional recovery after spinal cord injury. Cell Metab 31: 623

Han SM, Baig HS, Hammarlund M (2016) Mitochondria localize to injured axons to support regeneration. Neuron 92: 1308

Haraldsen JD, Liu G, Botting CH, Walton JGA, Storm J, Phalen TJ, Kwok LY, Soldati-Favre D, Heintz NH, Müller S, et al (2009) Identification of conoidin A as a covalent inhibitor of peroxiredoxin II. Org Biomol Chem 7: 3040–3048

Heiden MGV, Cantley LC, Thompson CB (2009) Understanding the Warburg effect: the metabolic requirements of cell proliferation. Science (1979) 324: 1029–1033

Hill CS, Coleman MP, Menon DK (2016) Traumatic axonal injury: mechanisms and translational opportunities. Trends Neurosci 39: 311–324

Ho J, Tumkaya T, Aryal S, Choi H, Claridge-Chang A (2019) Moving beyond P values: data analysis with estimation graphics. Nat Methods 2019 16:7 16: 565–566

Hopkins EL, Gu W, Kobe B, Coleman MP (2021) A novel NAD signaling mechanism in axon degeneration and its relationship to innate immunity. Front Mol Biosci 8: 703532

Van houcke J, Bollaerts I, Geeraerts E, Davis B, Beckers A, Van Hove I, Lemmens K, De Groef L, Moons L (2017a) Successful optic nerve regeneration in the senescent zebrafish despite age-related decline of cell intrinsic and extrinsic response processes. Neurobiol Aging 60: 1–10

Huang N, Li S, Xie Y, Han Q, Xu XM, Sheng ZH (2021) Reprogramming an energetic AKT-PAK5 axis boosts axon energy supply and facilitates neuron survival and regeneration after injury and ischemia. Curr Biol 31: 3098

Iwata R, Casimir P, Erkol E, Boubakar L, Planque M, Gallego López IM, Ditkowska M, Gaspariunaite V, Beckers S, Remans D, et al (2023) Mitochondria metabolism sets the species-specific tempo of neuronal development. Science (1979) 379

Jacobi A, Tran NM, Yan W, Benhar I, Tian F, Schaffer R, He Z, Sanes JR (2022) Overlapping transcriptional programs promote survival and axonal regeneration of injured retinal ganglion cells. Neuron 110: 2625-2645.e7

Ketschek A, Sainath R, Holland S, Gallo G (2021) The axonal glycolytic pathway contributes to sensory axon extension and growth cone dynamics. J Neurosci 41: 6637–6651

Kölsch Y, Hahn J, Sappington A, Stemmer M, Fernandes AM, Helmbrecht TO, Lele S, Butrus S, Laurell E, Arnold-Ammer I, et al (2021) Molecular classification of zebrafish retinal ganglion cells links genes to cell types to behavior. Neuron 109: 645-662.e9

Korade Z, Kenworthy AK (2008) Lipid rafts, cholesterol, and the brain. Neuropharmacology 55: 1265

Kramer AA, Olson GM, Chakraborty A, Blackmore MG (2021) Promotion of corticospinal tract growth by KLF6 requires an injury stimulus and occurs within four weeks of treatment. Exp Neurol 339: 113644

Kulkarni RA, Worth AJ, Zengeya TT, Shrimp JH, Garlick JM, Roberts AM, Montgomery DC, Sourbier C, Gibbs BK, Mesaros C, et al (2017) Discovering targets of non-enzymatic acylation by thioester reactivity profiling. Cell Chem Biol 24: 231

Lee-Liu D, Edwards-Faret G, Tapia VS, Larraín J (2013) Spinal cord regeneration: Lessons for mammals from non-mammalian vertebrates. Genesis 51: 529–544

Leibinger M, Andreadaki A, Diekmann H, Fischer D (2013) Neuronal STAT3 activation is essential for CNTF- and inflammatory stimulation-induced CNS axon regeneration. Cell Death Dis 2013 4:9 4: e805–e805

Li H, Guglielmetti C, Sei YJ, Zilberter M, Le Page LM, Shields L, Yang J, Nguyen K, Tiret B, Gao X, et al (2023) Neurons require glucose uptake and glycolysis in vivo. Cell Rep 42: 112335

Li H, Kuwajima T, Oakley D, Nikulina E, Hou J, Yang WS, Lowry ER, Lamas NJ, Amoroso MW, Croft GF, et al (2016) protein prenylation constitutes an endogenous brake on axonal growth. Cell Rep 16: 545–558

Liberti M V, Locasale JW (2016) The Warburg effect: how does it benefit cancer cells? Trends Biochem Sci 41: 211–218

Loreto A, Di Stefano M, Gering M, Conforti L (2015) Wallerian degeneration is executed by an NMN-SARM1-dependent late Ca2+ influx but only modestly influenced by mitochondria. Cell Rep 13: 2539–2552

Love MI, Huber W, Anders S (2014) Moderated estimation of fold change and dispersion for RNA-seq data with DESeq2. Genome Biol 15: 1–21

Luo X, Ribeiro M, Bray ER, Lee DH, Yungher BJ, Mehta ST, Thakor KA, Diaz F, Lee JK, Moraes CT, et al (2016) Enhanced transcriptional activity and mitochondrial localization of STAT3 co-induce axon regrowth in the adult central nervous system. Cell Rep 15: 398–410

Mar FM, Simões AR, Leite S, Morgado MM, Santos TE, Rodrigo IS, Teixeira CA, Misgeld T, Sousa MM (2014) CNS axons globally increase axonal transport after peripheral conditioning. J Neurosci 34: 5965

Masin L (2021) Neural Circuit Development and Regeneration / tectal-reinnervation-zebrafish · GitLab.

Masin L, Bergmans S, Van Dyck A, Farrow K, De Groef L, Moons L (2024) Local glycolysis supports injury-induced axonal regeneration. J Cell Biol 223

Masin L, Claes M, Bergmans S, Cools L, Andries L, Davis BM, Moons L, De Groef L (2021) A novel retinal ganglion cell quantification tool based on deep learning. Sci Rep 2021 11:1 11: 1–13

Mason MRJ, Van Erp S, Wolzak K, Behrens A, Raivich G, Verhaagen J (2021) The Jun-dependent axon regeneration gene program: Jun promotes regeneration over plasticity. Hum Mol Genet 31: 1242

Munemasa Y, Seok HK, Jae HA, Kwong JMK, Caprioli J, Piri N (2008) Protective effect of thioredoxins 1 and 2 in retinal ganglion cells after optic nerve transection and oxidative stress. Invest Ophthalmol Vis Sci 49: 3535–3543

OpenCV OpenCV - Open Computer Vision Library. https://opencv.org

Pajak B, Siwiak E, Sołtyka M, Priebe A, Zieliński R, Fokt I, Ziemniak M, Jaśkiewicz A, Borowski R, Domoradzki T, et al (2019) 2-Deoxy-d-Glucose and its analogs: from diagnostic to therapeutic agents. Int J Mol Sci 21: 234

Pedregosa F F, Michel V, Grisel O O, Blondel M, Prettenhofer P, Weiss R, Vanderplas J, Cournapeau D, Pedregosa F, Varoquaux G, et al (2011) Scikit-learn: Machine learning in Python. J Mach Learn Res 12: 2825–2830

Peregrín-Alvarez JM, Sanford C, Parkinson J (2009) The conservation and evolutionary modularity of metabolism. Genome Biol 10: 1–17

Petrova V, Nieuwenhuis B, Fawcett JW, Eva R (2021) Axonal organelles as molecular platforms for axon growth and regeneration after injury. Int J Mol Sci 22: 1–30

Poulain FE, Chien C Bin (2013) Proteoglycan-mediated axon degeneration corrects pretarget topographic sorting errors. Neuron 78: 49–56

Ramanathan RK, Kirkpatrick DL, Belani CP, Friedland D, Green SB, Chow HHS, Cordova CA, Stratton SP, Shadow ER, Baker A, et al (2007) A phase I pharmacokinetic and pharmacodynamic study of PX-12, a novel inhibitor of thioredoxin-1, in patients with advanced solid tumors. Clin Cancer Res 13: 2109–2114

Rhee SG, Kil IS (2017) Multiple functions and regulation of mammalian peroxiredoxins. Annu Rev Biochem 86: 749–775

Robinson BH, Petrova-Benedict R, Buncic JR, Wallace DC (1992) Nonviability of cells with oxidative defects in galactose medium: A screening test for affected patient fibroblasts. Biochem Med Metab Biol 48: 122–126

Roselló-Busquets C, de la Oliva N, Martínez-Mármol R, Hernaiz-Llorens M, Pascual M, Muhaisen A, Navarro X, del Valle J, Soriano E (2019) cholesterol depletion regulates axonal growth and enhances central and peripheral nerve regeneration. Front Cell Neurosci 13

Schaeffer J, Vilallongue N, Decourt C, Saudou F, Nawabi H, Correspondence SB, Blot B, Bakdouri N El, Plissonnier E, Excoffier B, et al (2023) Customization of the translational complex regulates mRNA-specific translation to control CNS regeneration. Neuron 111: 2881–2898.e12.

Schindelin J, Arganda-Carreras I, Frise E, Kaynig V, Longair M, Pietzsch T, Preibisch S, Rueden C, Saalfeld S, Schmid B, et al (2012) Fiji: an open-source platform for biological-image analysis. Nat Methods 2012 9:7 9: 676–682

Shabanzadeh AP, Charish J, Tassew NG, Farhani N, Feng J, Qin X, Sugita S, Mothe AJ, Wälchli T, Koeberle PD, et al (2021) Cholesterol synthesis inhibition promotes axonal regeneration in the injured central nervous system. Neurobiol Dis 150: 105259

Shigeoka T, Jung H, Jung J, Turner-Bridger B, Ohk J, Lin JQ, Amieux PS, Holt CE (2016) Dynamic axonal translation in developing and mature visual circuits. Cell 166: 181–192

van Spronsen M, Mikhaylova M, Lipka J, Schlager MA, van den Heuvel DJ, Kuijpers M, Wulf PS, Keijzer N, Demmers J, Kapitein LC, et al (2013) TRAK/Milton motor-adaptor proteins steer mitochondrial trafficking to axons and dendrites. Neuron 77: 485–502

Sun F, Park KK, Belin S, Wang D, Lu T, Chen G, Zhang K, Yeung C, Feng G, Yankner BA, et al (2011) Sustained axon regeneration induced by co-deletion of PTEN and SOCS3. Nature 480: 372–375

Szwed A, Kim E, Jacinto E (2021) regulation and metabolic functions of mTORC1 and mTORC2. Physiol Rev 101: 1371–1426

Tang B (2022) Cholesterol synthesis inhibition or depletion in axon regeneration. Neural Regen Res 17: 271–276

Tapia ML, Nascimento-dos-Santos G, Park KK (2022) Subtype-specific survival and regeneration of retinal ganglion cells in response to injury. Front Cell Dev Biol 10: 956279

TeSlaa T, Ralser M, Fan J, Rabinowitz JD (2023) The pentose phosphate pathway in health and disease. Nature Metabolism 2023 5:8 5: 1275–1289

Tonge D, Chan K, Zhu N, Panjwani A, Arno M, Lynham S, Ward M, Snape A, Pizzey J (2008) Enhancement of axonal regeneration by in vitro conditioning and its inhibition by cyclopentenone prostaglandins. J Cell Sci 121: 2565–2577

Varadarajan SG, Hunyara JL, Hamilton NR, Kolodkin AL, Huberman AD (2022) Central nervous system regeneration. Cell 185: 77–94

Van Der Walt S, Schönberger JL, Nunez-Iglesias J, Boulogne F, Warner JD, Yager N, Gouillart E, Yu T (2014) Scikit-image: Image processing in python. PeerJ 2014: e453

Wang H, Wu M, Zhan C, Ma E, Yang M, Yang X, Li Y (2012) Neurofilament proteins in axonal regeneration and neurodegenerative diseases. Neural Regen Res 7: 620–626

Wang M, Casey PJ (2016) Protein prenylation: unique fats make their mark on biology. Nat Rev Mol Cell Biol 2016 17:2 17: 110–122

Wang X, Ma C, Nie L (2022) GAP-43 induces the differentiation of bone marrow-derived mesenchymal stem cells into retinal ganglial-like cells. Comput Math Methods Med 2022: 1–7

Williams PR, Benowitz LI, Goldberg JL, He Z (2020) Axon regeneration in the mammalian optic nerve. Annu Rev Neurosci 43: 473–494.

Wolfe AD, Koberstein JN, Smith CB, Stewart ML, Gonzalez IJ, Hammarlund M, Hyman AA, Stork PJS, Goodman RH, Colón-Ramos DA (2024) Local and dynamic regulation of neuronal glycolysis in vivo. Proc Natl Acad Sci U S A 121: e2314699121

Xu Y, Chen M, Hu B, Huang R, Hu B (2017) In vivo imaging of mitochondrial transport in single-axon regeneration of zebrafish mauthner cells. Front Cell Neurosci 11: 4

Zhou B, Yu P, Lin MY, Sun T, Chen Y, Sheng ZH (2016) Facilitation of axon regeneration by enhancing mitochondrial transport and rescuing energy deficits. J Cell Biol 214: 103

Zorov DB, Juhaszova M, Sollott SJ (2014) Mitochondrial reactive oxygen species (ROS) and ROS-induced ROS release. Physiol Rev 94: 909–950

