## Supplementary figures and images for "Successful axonal regeneration is driven by evolutionarily conserved metabolic reprogramming"

### Supplemental figure 01

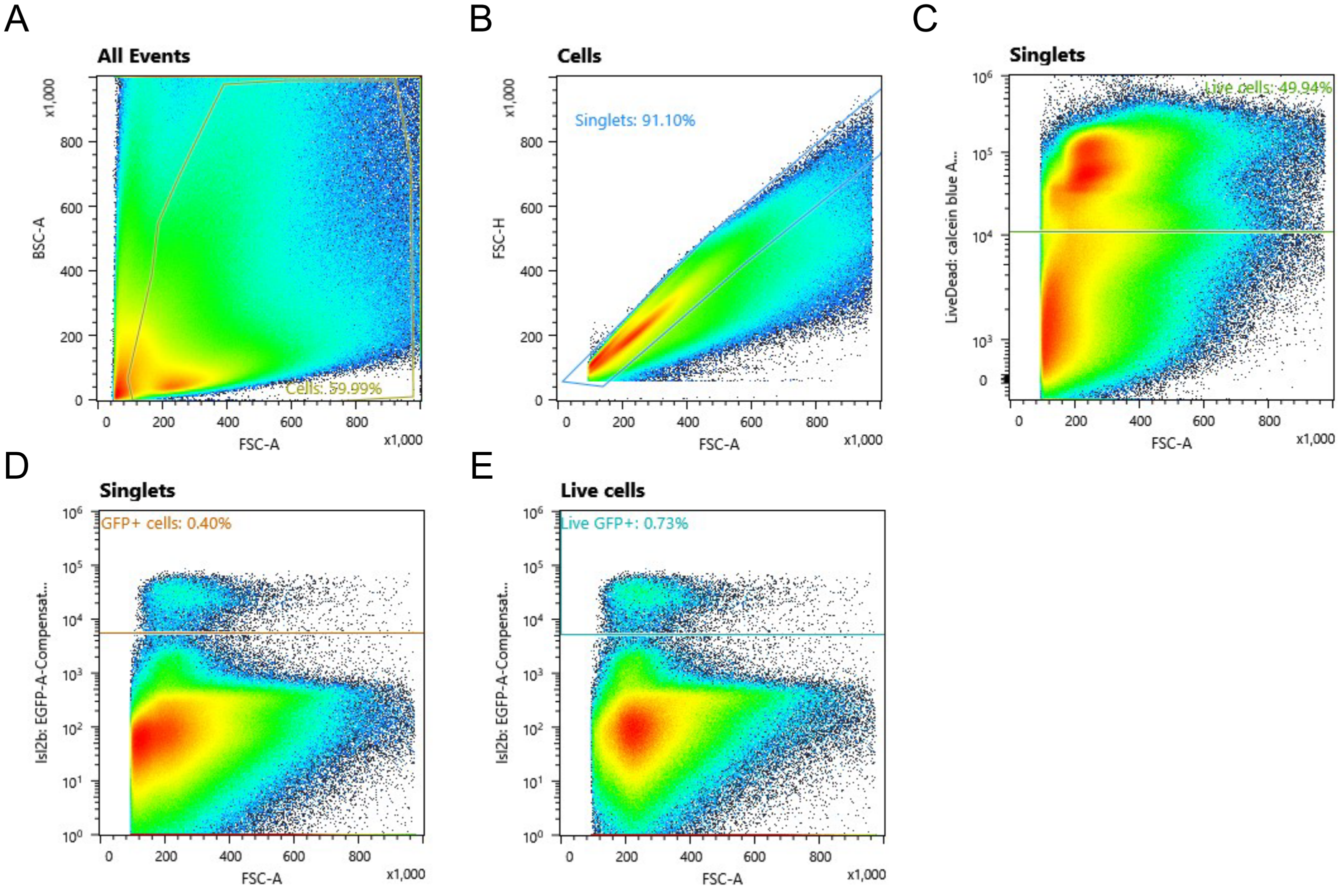

### Supplemental figure 02

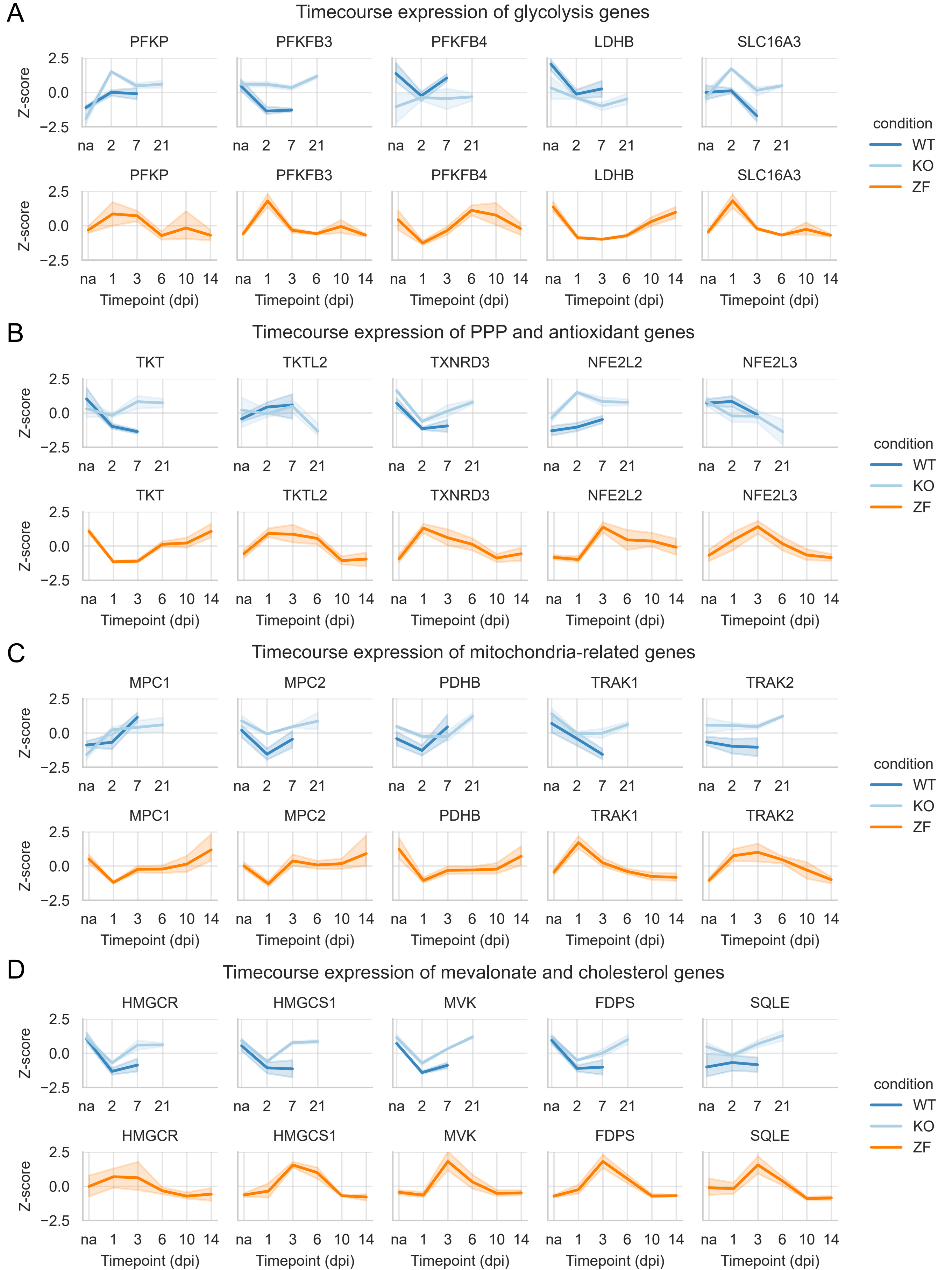

### Supplemental figure 03

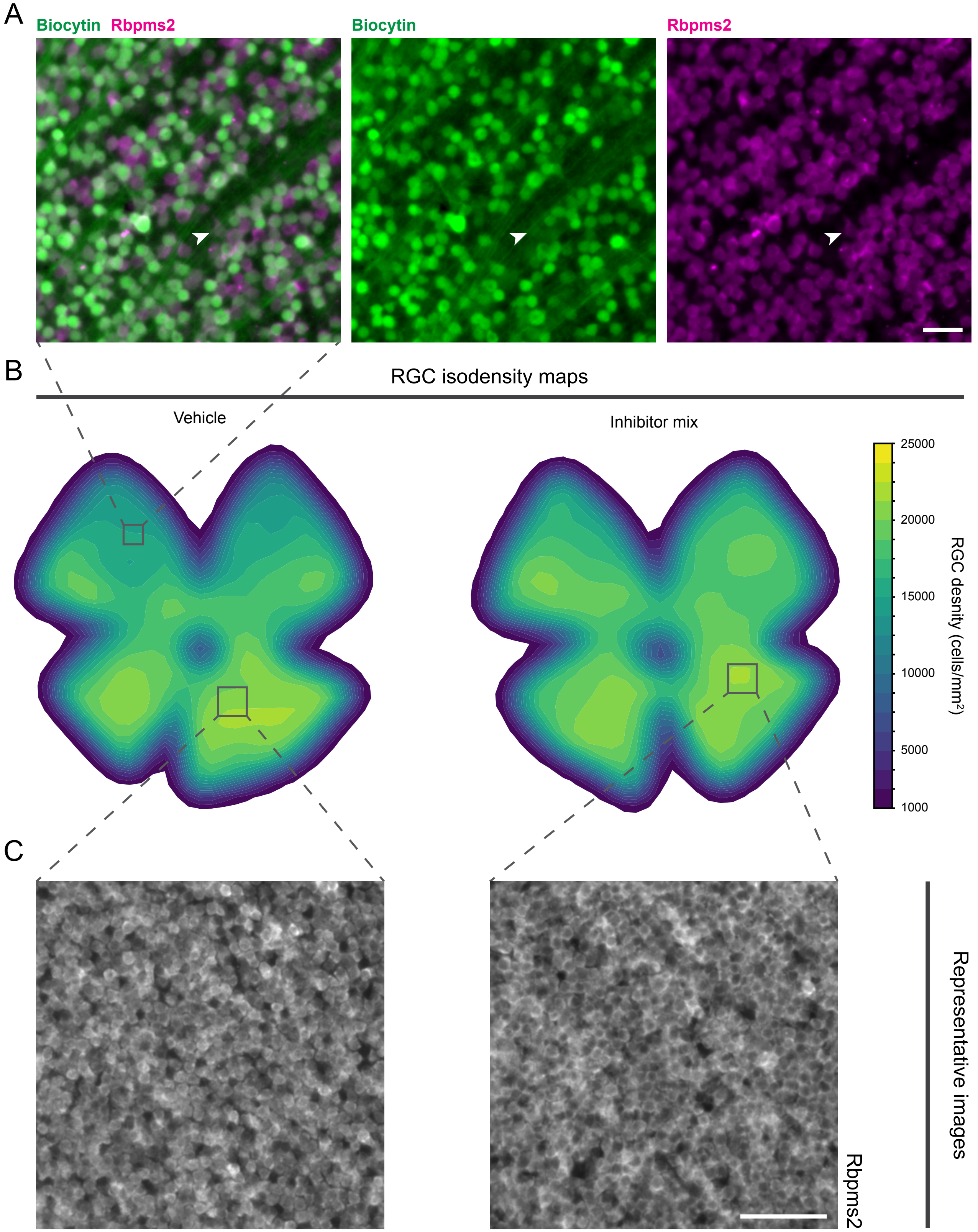

### Supplemental figure 04

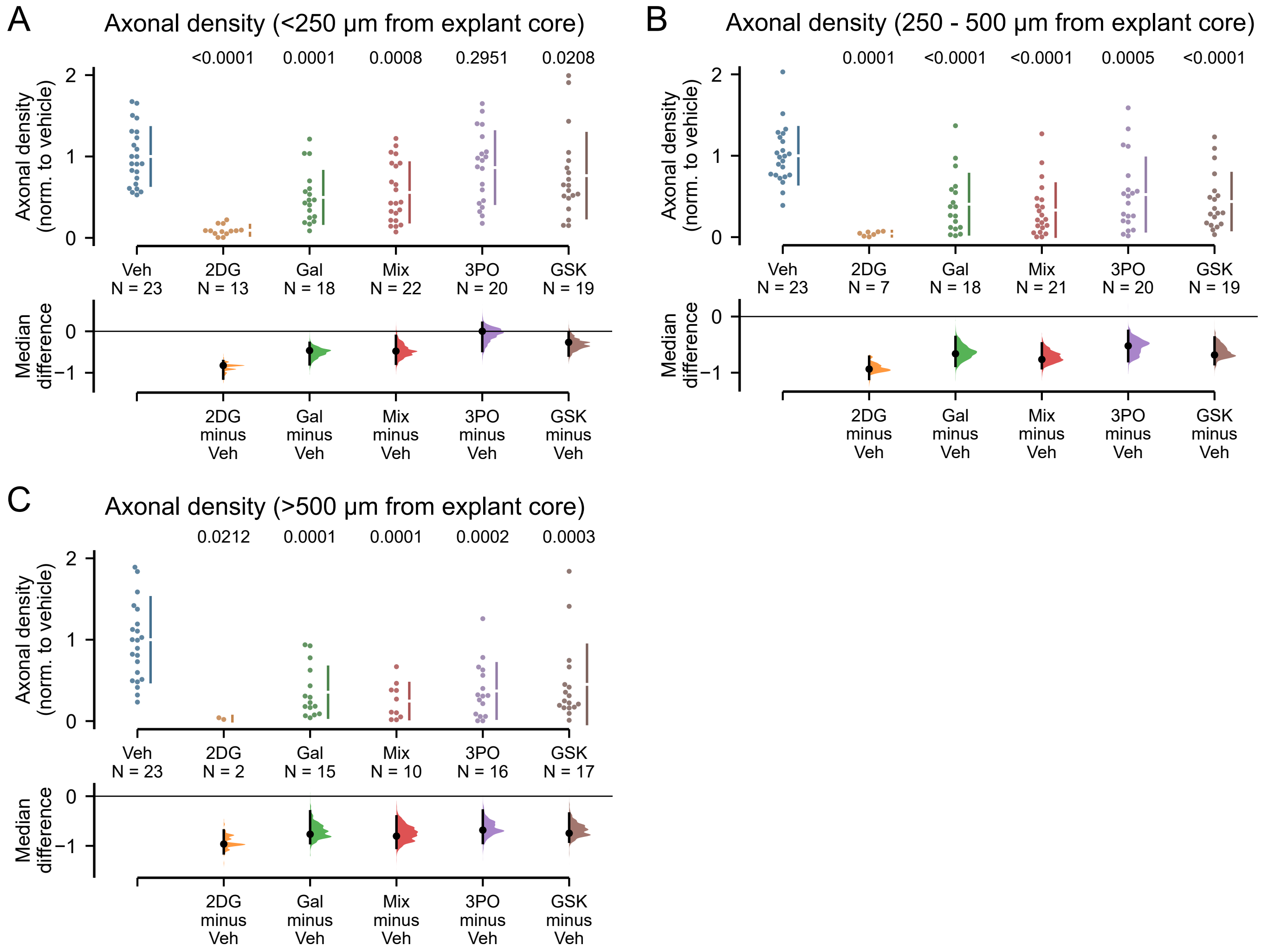
